# Recombination persists near the mating-type locus in pseudo-homothallic *Cladorrhinum* fungi (Sordariales)

**DOI:** 10.64898/2026.10.08.757285

**Authors:** Jeanne Legendre–Despas, Emy Cavicchi, Valérie Gautier, Christophe Lalanne, Tatiana Giraud, Philippe Silar, Fanny E. Hartmann

**Affiliations:** Écologie, Société et Évolution, CNRS, Université Paris-Saclay, AgroParisTech, Gif-sur-Yvette, France; Laboratoire Interdisciplinaire des Énergies de Demain, Université Paris Cité, Paris, France

**Keywords:** *Podospora australis*, *Podospora anserina*, sex chromosomes

## Abstract

Recombination suppression has evolved repeatedly across eukaryotes around sex-determining and mating-type loci. Recent theoretical models proposed that such suppression can evolve due to various effects of recessive deleterious mutations, predicting that it should only evolve in lineages experiencing extended diploid phases. Ascomycete fungi provide a powerful system to test this prediction, as pseudo-homothallic ascomycetes maintain prolonged diploid-like phases, while most other ascomycetes have a predominant haploid phase. Megabase-scale non-recombining regions have in fact been reported around the mating-type locus in the four pseudo-homothallic ascomycete lineages studied to date. Here, we studied a previously unexplored lineage of *Sordariales*, formerly referred to as *Podospora australis* and suggested to be pseudo-homothallic. Combining genome sequencing, phylogenomic analyses and life-cycle characterization, we identified and described two distinct species and reassigned them to the *Cladorrhinum* genus. We showed that the two species produce large proportions of self-fertile ascospore-derived mycelia, but the frequencies varied substantially among four strains, and with a selfing progeny, indicating large variation in the extent of the diploid-like phase. By analysing the genomes of five wild strains and of selfing progenies, we did not detect any footprints of large recombination suppression (no heterozygosity, rearrangements or repeated elements). This may be due to the smaller proportion of the life cycle spent in a diploid-like stage compared to other pseudo-homothallic species, or to other features of the life cycle, in agreement with theoretical models. In addition, our results decouple the evolution of pseudo-homothallism from that of recombination suppression, showing that there is no fundamental developmental constraint forcing their joint evolution.

## Introduction

By reshuffling alleles into new combinations, recombination increases genetic diversity, facilitates the purging of deleterious mutations, and enhances the efficacy of natural selection (Muller 1932; Hill and Robertson 1966). Despite its advantages, recombination is often suppressed in specific genomic regions. In particular, recombination suppression has evolved repeatedly around sex-determining loci in many animals and plants, forming sex chromosomes (Charlesworth 2017; Furman et al. 2020; Saunders and Muyle 2024). In fungi too, recombination suppression has evolved around mating-type loci independently in multiple lineages (Hartmann et al. 2021b; Duhamel et al. 2022; Jay et al. 2024; De Filippo et al. 2026). In non-recombining regions, chromosomes carrying alternative sex-determining or mating-type alleles cease exchanging genetic material and evolve independently, resulting in sequence divergence and degeneration due to the reduced efficacy of selection (Muller 1932; Hill and Robertson 1966; Smith and Haigh 1974).

The evolutionary causes of recombination suppression remain actively debated. For decades, the dominant explanation has been the sexually antagonistic selection hypothesis, which proposes that recombination suppression is favored between a sex-determining locus and alleles that benefit one sex but are detrimental to the other (Rice 1984). However, there has been very little support, if any, for this hypothesis (Ironside 2010; Jay et al. 2024). Furthermore, this hypothesis cannot apply to the fungal mating-type chromosomes, as there is no known antagonistic selection between mating types (Bazzicalupo et al. 2019). Furthermore, the ascomycete fungi with suppressed recombination are precisely the rare ones whose life cycle lacks a haploid phase, which is the phase where antagonistic selection could act, i.e., in which individual cells are normally characterized by one mating type or the other (Ironside 2010; Hartmann et al. 2021b; Jay et al. 2024).

Alternative hypotheses have recently gained attention, including some based on neutral accumulation of divergence (Jeffries et al. 2021), or proposing that recombination suppression could evolve because of the presence of recessive deleterious mutations in genomes (Charlesworth and Wall 1999; Antonovics and Abrams 2004; Lenormand and Roze 2022; Jay et al. 2024; Jay et al. 2025; Guyot et al. 2026). Some of these latter models propose that inversions capturing fewer deleterious mutations than average would have a selective advantage. These inversions could reach fixation even if they still carry some recessive deleterious mutations, provided they also capture an allele that is maintained in a permanently heterozygous state, e.g., a sex-determining locus or a mating-type locus. Indeed, the recessive deleterious mutations would then be sheltered as permanently heterozygous (Lenormand and Roze 2022; Jay et al. 2024; Jay et al. 2025). In another recent model, pseudo-overdominance could evolve in the flanking regions of a mating-type locus, i.e., with recessive deleterious mutations in repulsion, under specific conditions depending on parameters such as recombination and mutation rates (Guyot et al. 2026). This genetic architecture could make recombination deleterious and thereby favor the evolution of recombination suppression (Guyot et al. 2026).

A key prediction of these models based on recessive deleterious mutations is that recombination suppression around permanently heterozygous loci, such as sex-determining loci or mating-type loci, should only evolve in organisms with a predominantly diploid life cycle, because only then can recessive deleterious alleles segregate in populations, and remain masked from selection in heterozygotes. Testing this prediction requires comparative systems displaying variation in the duration and importance of diploid stages during the life cycle. Fungi provide particularly powerful systems for investigating this question. Indeed, fungal life cycles vary considerably in the relative importance of haploid and diploid-like phases and some have recombination suppression on their mating-type chromosomes, providing a unique opportunity to investigate the relationship between prolonged diploid-like phases and the evolution of recombination suppression (Jay et al. 2024; De Filippo et al. 2026). In addition, they have short generation times, are experimentally tractable, and possess compact genomes, making them important models in evolutionary genomics (Gladieux et al. 2014).

Among fungi, the taxonomically diverse *Sordariales* order (Ascomycota) is especially relevant because pseudo-homothallism has evolved independently in several lineages. This reproductive strategy involves a prolonged heterokaryotic phase during which genetically distinct and unfused haploid nuclei of opposite mating types coexist, often with more than two nuclei within the same hyphal compartments. Such a diploid-like life cycle allows recessive deleterious mutations to remain sheltered around the permanently heterozygous mating-type locus, contrasting with the strictly haploid life cycle found in most Sordariales, producing spores and mycelia carrying a single type of haploid nuclei. According to the theoretical models based on the effects of recessive deleterious mutations, one would therefore expect that recombination suppression only evolves in pseudo-homothallic Sordariales species, although not necessarily in all of them, as other parameters can be important for the evolution of recombination suppression in these models (Jay et al. 2025; Guyot et al. 2026).

In Sordariales, as is generally the case in ascomycetes, the two mating types are determined by two highly differentiated sequences at the mating-type (MAT) locus. These sequences differ in both the number and identity of the genes they contain and are therefore referred to as idiomorphs rather than alleles. They are highly conserved across *Pezizomycotina* fungi and are called MAT1-1 and MAT1-2 (Butler 2007; Debuchy et al. 2010; Dyer et al. 2016). In the haploid-like ascomycetes, karyogamy between nuclei of opposite mating types occurs within the developing ascus and is immediately followed by meiosis, producing four haploid nuclei; in many species, a subsequent post-meiotic mitosis results in eight haploid nuclei. Each nucleus is then individually packaged into an ascospore, resulting in eight uni-nucleate spores per ascus. In contrast, the pseudo-homothallic *Sordariales* fungi package pairs of nuclei of opposite mating types into the same ascospore, producing four large heterokaryotic spores (Figure S1A). These heterokaryotic spores, i.e., carrying both mating types, germinate into self-fertile mycelia capable of completing the sexual cycle without a distinct mating partner. This is actually why they are named pseudo-homothallic, as homothallic fungi can also mate within the same thallus, but the mycelium being haploid (Debuchy et al. 2010; Grognet and Silar 2015).

Pseudo-homothallic *Sordariales* fungi also provide a particularly suitable system for identifying regions of recombination suppression. Because reproduction occurs predominantly through selfing between nuclei of opposite mating types originating from the same ascospore, i.e. diploid selfing (Billiard et al. 2012), most of the genome is highly homozygous. In contrast, when recombination is suppressed around the MAT locus, regions linked to the MAT locus accumulate sequence divergence and remain heterozygous, as the MAT locus is obligatorily heterozygous. These regions therefore appear as large blocks of heterozygosity surrounded by an otherwise homozygous genome (Hartmann et al. 2021a,b; Vittorelli et al. 2023; Silar et al. 2025; De Filippo et al. 2026). Heterozygosity around the MAT locus and in some localised stretches elsewhere in genomes may also be due to occasional outcrossing events that still sometimes occur in these fungi. Genuine recombination suppression can be distinguished from residual heterozygosity resulting from recent outcrossing events by following their persistence through successive generations of selfing. Heterozygosity persisting around the MAT locus for longer than expected given the number of selfing generations and the local recombination rate is indicative of recombination suppression (Hartmann et al. 2021b).

Megabase-scale recombination suppression has been documented around the MAT locus in all pseudo-homothallic *Sordariales* studied to date, including all members of the species complexes of *Podospora anserina* (family *Podosporaceae;* (Grognet et al. 2014; Hartmann et al. 2021a; Silar et al. 2025)), *Neurospora tetrasperma* (family *Sordariaceae* (Menkis et al. 2008; Sun et al. 2017)), *Schizothecium tetrasporum* and *Schizothecium pseudotetrasporum* (family *Schizotheciaceae;* (Vittorelli et al. 2023; Guyot et al. 2025; De Filippo et al. 2026)). In these species thriving in soil and herbivore dung, heterokaryotic ascospores are produced at very high frequencies (>99%) and recombination suppression extends over large genomic regions surrounding the MAT locus (>0.6 Mb). The proximal mechanisms allowing the production of heterokaryotic ascospores are different across these lineages. In *N. tetrasperma*, the MAT locus is fully linked to the centromere, causing first-division segregation; together with overlapping meiotic spindles, this ensures that nuclei of opposite mating types are packaged into the same ascospore (Raju and Perkins 1994; Menkis et al. 2008). In contrast, in *P. anserina* and in *S. tetrasporum* sensu stricto, an obligate crossover between the MAT locus and the centromere (>1 Mb apart) causes second-division segregation, which, through a different spindle arrangement, also results in the packaging of opposite mating types within each ascospore ((Figure S1A), (Raju and Perkins 1994; Grognet et al. 2014; Vittorelli et al. 2023)). In very rare meioses, no cross-over occurs, or a double cross-over, which leads to first-division segregation of the mating-type locus at meiosis, and to the packaging of two nuclei of the same mating type within each ascospore (Marcou et al. 1979; Vittorelli et al. 2023). In addition, the heterokaryotic mycelium sometimes loses nuclei of one mating type or displays desequilibria between nuclei of different mating types (Figure S1B), as experimentally shown in the *S. tetrasporum* species complex (Vittorelli et al. 2023; Guyot et al. 2025). For example, while 13% of ascospore-derived mycelia was found homokaryotic based on PCR genotyping in the CBS815.71 *S. tetrasporum* s.s. strain, mycelia derived from other ascospores of the same asci were heterokaryotic, suggesting second-division segregation of the mating-type locus at meiosis; such second-division segregation would imply that all four ascospores are initially heterokaryotic and that growing mycelia subsequently lost nuclei of one mating type (Vittorelli et al. 2023).

The recombination suppression between the MAT locus and the centromere in *N. tetrasperma* thus likely plays a role in the development of heterokaryotic spores, but recombination suppression also progressively extended farther away from the MAT locus (Menkis et al. 2008; Hartmann et al. 2021b). In *P. anserina* and *Schizothecium* pseudohomothallic species complexes, the recombination suppression is thought to play no role in the packaging of nuclei of opposite mating types in ascospores (Vittorelli et al. 2023), and recombination suppression also extended progressively away from the mating-type locus in multiple species of the species complexes (Hartmann et al. 2021a; Vittorelli et al. 2023; De Filippo et al. 2026).

Four independent events of joint evolution of recombination suppression and pseudo-homothallism have been inferred to date (Jay et al. 2024; De Filippo et al. 2026). In contrast, no recombination suppression has been reported in *Sordariales* fungi with a prominent homokaryotic life cycle (Jay et al. 2024; De Filippo et al. 2026). In this context, we investigated a previously unexplored *Sordariales* lineage, the species previously known as *Podospora australis* found in dung. This species is often, but not always, synonymized with *Podospora taenioides* (Mirza and Cain 1969; Lundqvist 1972; Doveri 2004). Both species produce similar-looking four-spored asci and have been suggested to be pseudo-homothallic in the literature. This lineage has indeed been described as producing four-spored asci, as typical in pseudo-homothallic ascomycetes (Dodge 1936; Mainwaring 1966). However, it has been little studied and it remains unknown whether this species is truly pseudo-homothallic, i.e. whether it produces heterokaryotic spores containing nuclei of opposite mating types that germinate into self-fertile mycelia. The meiotic segregation pattern at the MAT locus and the presence of recombination suppression are also unknown.

We therefore studied the lineage currently referred to as *P. australis* or *P. taenioides,* in order to clarify its taxonomy, to check whether it was genuinely pseudo-homothallic and to test whether recombination was suppressed around its MAT locus. To this end, we isolated nine wild strains belonging to this lineage from herbivore dung in France and Italy. As cryptic species are often detected within morphological species in *Sordariales* fungi (Boucher et al. 2017; Gladieux et al. 2020; De Filippo et al. 2025; Silar et al. 2025), we used whole genome sequences to delimit two distinct species based on their average nucleotide divergence (ANI), and studied their morphological specificities. We assigned them to the *Cladorrhinum* genus based on phylogenetic analyses and provided a description of both species, *C. taenioides* and *C. neoaustrale*. We then quantified the frequency of ascospore-derived heterokaryotic mycelia based on MAT locus genotyping and self-fertility. We also generated a high-quality reference genome based on long-read sequencing, and examined patterns of genomic heterozygosity around the MAT locus in five wild isolates, as well as in selfed progeny. We found that both species produced high proportions of heterokaryotic ascospore-derived mycelia, although this proportion varied among strains, suggesting variation in the extent of the diploid-like phase. We detected no evidence of extensive recombination suppression, contrasting with the megabase-scale non-recombining regions reported in other pseudo-homothallic Sordariales. By expanding the taxonomic breadth of studied pseudo-homothallic *Sordariales*, our work contributes to our understanding of the relationship between the importance of the diploid-like phase and the evolution of recombination suppression around mating-type loci in fungi.

## Results

### Two distinct species assigned to the *Cladorrhinum* genus with self-fertile ascospore-derived mycelium

Nine fungal strains with morphological features similar to that of *P. australis* (Speg.) Niessl and *P. taenioides* (Griffiths) Cain (Mirza and Cain 1969) were recovered from herbivore dung (Table 1, see Taxonomy section for morphological description). As a first molecular characterization, their internal transcribed spacers (ITS) were sequenced, which revealed variation at two positions. On the one hand, the strains PSN303, PSN309, PSN314, PSN873 and PSN1235 had a C at position 56 and two As after position 136, the position 1 corresponding to the first nucleotide after the end of the ITS1 primer; on the other hand, the strains PSN672, PSN824, PSN993 and PSN1024 had a T at position 56 and three As after position 136. This low polymorphism level suggested that all nine strains belonged to a single species or to two closely related species. When the ITS sequences were used to search with BLAST the GenBank RefSeq database, compiling the ITS sequence of all type strains (https://www.ncbi.nlm.nih.gov/bioproject/PRJNA177353/), the closest hits corresponded to various *Cladorrhinum* species (as of May 14^th^, 2026, the best hit was with *Cladorrhinum queenslandicum* BRIP 63076a). This confirmed that the strains belonged to the *Sordariales* order, and more specifically to the *Podosporaceae* family. When the ITS sequences were used to search the general database, the best hits corresponded to several strains of *P. australis*, including strains TTI-313 and TTI-248, which have been shown to produce interesting secondary metabolites (Li et al. 2016). Note that the strain labelled as *P. australis* commercially available in the Westerdijk Institute collection, CBS 370.59, does not have its ITS region sequenced. Only its partial LSU sequence (GenBank: MH869435.1) is currently available, and is nearly identical to that of *Didymella prosopidis* CBS 136414, a species belonging to the *Dothideomycetes*, suggesting that CBS370.59 actually does not belong to *P. australis*. All nine strains isolated from single ascospores initially produced perithecia, i.e. the sexual structures of Sordariales carrying ascospores resulting from meiosis, indicating that they were self-fertile and therefore pseudo-homothallic. Some of them, however, stopped producing perithecia after being replated on Petri dishes, which could be due to the loss of nuclei of one mating type during mycelial growth, as observed in the *S. tetrasporum* species complex (Vittorelli et al. 2023; Guyot et al. 2025).

**Table 1.** Characteristics of the studied *Cladorrhinum* strains: strain ID, species name, and information on the collection year, geographical location and substrate.

| Strain ID | Species | Isolation year | Geographical location | Substrate |
| --- | --- | --- | --- | --- |
| PSN303 | *C. taenioides* | 2018 | Cassimorenga, Apennines, Italy | Rabbit dung |
| PSN309 | *C. taenioides* | 2018 | Cassimorenga, Apennines, Italy | Rabbit dung |
| PSN314 | *C. taenioides* | 2018 | Cassimorenga, Apennines, Italy | Rabbit dung |
| PSN672 | *C. neoaustrale* | 2021 | Camargue, France | Horse dung |
| PSN824 | *C. neoaustrale* | 2021 | Bertre, Tarn, France | Hare dung |
| PSN873 | *C. taenioides* | 2021 | Cassimorenga, Apennines, Italy | Horse dung |
| PSN993 | *C. neoaustrale* | 2021 | Camargue, France | Horse dung |
| PSN1024 | *C. neoaustrale* | 2021 | St Remy de Provence, France | Horse dung |
| PSN1235 | *C. taenioides* | 2022 | Montpellier, France | Cow dung |

We sequenced the genomes of the nine isolated strains using the short-read Illumina technology and assembled them *de novo.* DNA was either extracted from heterokaryotic mycelia or from the two homokaryotic mycelia of opposite mating types deriving from from the original heterokaryotic mycelia, either by spore isolation or mycelium fragmentation. Genome assembly sizes ranged from 37.0 Mb to 39.1 Mb (Table S1). The size differences were linked to variations in repeat content, likely due to different assembly procedures (Table S1). BUSCO scores were higher than 97.8%, showing that all assemblies were of good qualities. The MAT idiomorph identity of each genome was determined by blast analyses using the *P. anserina* MAT idiomorphs as queries. Notably, we detected a single MAT idiomorph in most of the genomes (Table S1), even those obtained from strains considered heterokaryons, based on the presence of perithecia in the initial fungal cultures. This further suggests that, during fungal culture prior to DNA extraction for sequencing, one mating-type nucleus may have been lost or that the relative abundance of nuclei of the two mating types may have become highly unbalanced (Figure S1B). For phylogeny analyses, we used a single genome assembly per strain (indicated with an asterisk in Table S1).

Average nucleotide identity (ANI) analyses indicated that the nine strains clustered into two distinct species, with ANI values above 99.8% among strains of the same species and approximately 97.7–97.8% between species. Examples of typical results obtained are presented in Figure 1 and all numerical values are reported in Table 2. The two distinct ITS sequences corresponded to the two species inferred with ANI. PSN303, PSN309, PSN314, PSN873 and PSN1235 belonged to one species, while PSN672, PSN824, PSN993 and PSN1024 belonged to the other species, referred hereafter as *C. taenioides* and *C. neoaustrale*, respectively (see Taxonomy section below).

**Figure 1.**
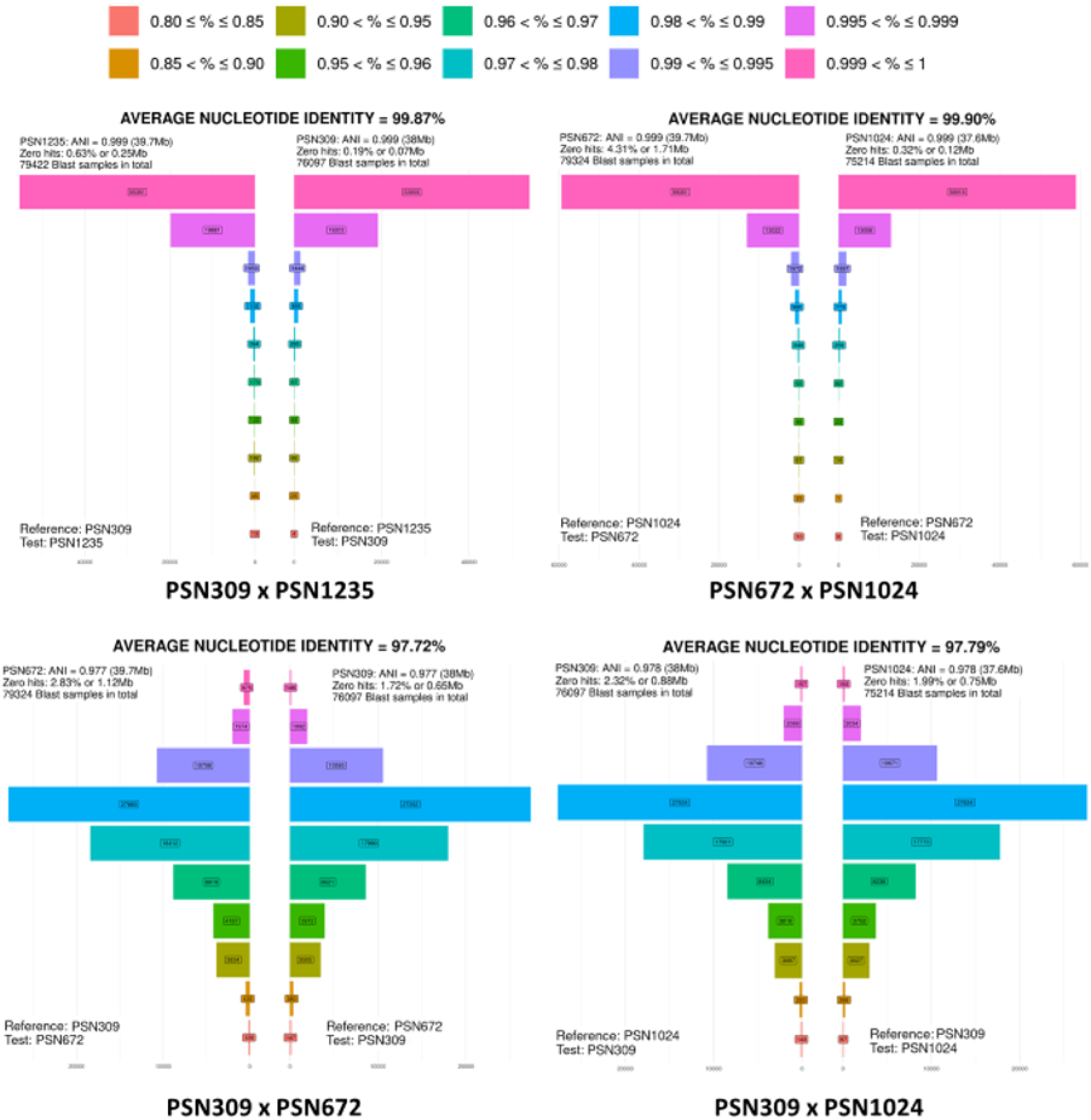
Representative graphics obtained using average nucleotide identity (ANI) calculation using FungANI. Horizontal bars quantify the number of BLAST results for each range of similarity percentages defined and color-coded at the top of the plots. The calculations were made in both directions, with on the left one strain as query (test) and the other as subject (reference) and reciprocally on the right. The final ANI is the average of the two values. The numbers of BLAST obtained for each percentage range are boxed in the corresponding bars and the total number of BLAST made during the analysis is given at the top of the plots. The zero hits correspond to BLAST results where no similarity was found. For more information see Reference (Lalanne and Silar 2025).

**Table 2.** Average nucleotide identities (ANI) of pairwise assembly comparisons (%). The used assemblies are labelled with an asterisk (*) in Table S1. ANI values between *Cladorrhinum neoaustrale and C. taenioides* are highlighted in bold.

|  |  | *C. taenioides* | *C. taenioides* | *C. taenioides* | *C. neoaustrale* | *C. neoaustrale* | *C. taenioides* | *C. neoaustrale* | *C. neoaustrale* |
| --- | --- | --- | --- | --- | --- | --- | --- | --- | --- |
|  |  | *PSN303* | *PSN309* | *PSN314* | *PSN672* | *PSN824* | *PSN873* | *PSN993* | *PSN1024* |
| *C. taenioides* | *PSN309* | 99.87 |  |  |  |  |  |  |  |
| *C. taenioides* | *PSN314* | 99.88 | 99.89 |  |  |  |  |  |  |
| *C. neoaustrale* | *PSN672* | **97.73** | **97.72** | **97.73** |  |  |  |  |  |
| *C. neoaustrale* | *PSN824* | **97.75** | **97.75** | **97.75** | 99.85 |  |  |  |  |
| *C. taenioides* | *PSN873* | 99.89 | 99.85 | 99.9 | **97.71** | **97.71** |  |  |  |
| *C. neoaustrale* | *PSN993* | **97.73** | **97.72** | **97.73** | 99.88 | 99.8 | **97.73** |  |  |
| *C. neoaustrale* | *PSN1024* | **97.78** | **97.79** | **97.77** | 99.90 | 99.88 | **97.78** | 99.94 |  |
| *C. taenioides* | *PSN1235* | 99.89 | 99.87 | 99.88 | **97.7** | **97.71** | 99.96 | **97.73** | **97.77** |

The phylogeny based on ITS + LSU sequences placed the nine strains in the *Cladorrhinum* genus within the *Podosporaceae* family (Figure 2). The dataset included *Podosporaceae* species with available genomes, as well as *Cladorrhinum* and *Stchigelomyces* species for which both ITS and LSU regions were available in GenBank (Table S2). The tree was rooted with *Similihumicola thailandica*, which belongs to the family most closely related to the *Podosporaceae* (Tangthirasunun et al. 2026). All nine strains clustered together in the genus *Cladorrhinum* close to *C. queenslandicum*, as expected from the RefSeq BLAST results. The two species identified by the ANI analysis were clearly separated, and each received strong bootstrap support (95%). The closest heterothallic species are likely *C. foecundissimum* and *C. grandiusculum* (P. Silar, personal com).

**Figure 2.**
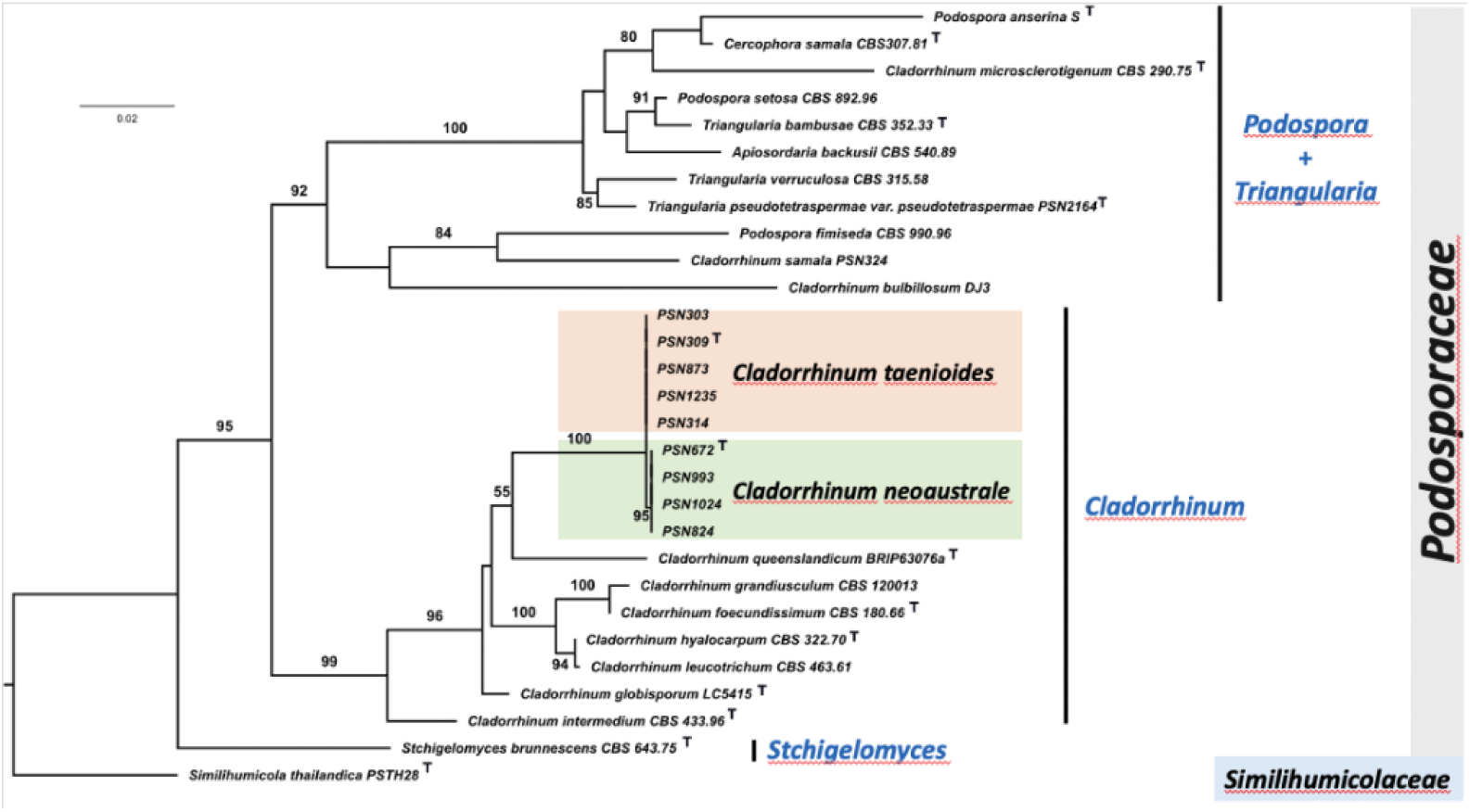
Phylogram of selected *Podosporaceae* species generated with IQ-tree from maximum likelihood analysis based on ITS + LSU sequences. Nineteen strains were included in the combined sequence analysis, which comprised 1153 characters with gaps; number of constant sites: 885; number of invariant (constant or ambiguous constant) sites: 885; number of parsimony informative sites: 174; number of distinct site patterns: 314. The tree was rooted with *Similihumicola thailandica*. The log-likelihood of the tree was - 4527.0058. Only ultrafast bootstrap values higher than 80% are displayed. The ^T^ symbol indicates the type specimen. The two focal *Cladorrhinum* species are highlighted in orange and green. The known genera of the *Podosporaceae* are in blue.

In order to obtain a better phylogenetic resolution and accurately place the *Cladorrhinum* genus in the phylogeny of Podosporaceae, we performed a phylogenomic analysis, which also grouped the two *Cladorrhinum* species together and separated with 100% statistical support the two species (Figure 3; Table S3). They branched as a sister group to the *Triangularia* genus defined by (Wang et al. 2019), with 100% statistical support, the *Podospora* genus as defined by Wang et al. being their closest outgroup (Figure 3). Note that our ITS + LSU analyses did not place the genus at this position (Figure 2).

**Figure 3.**
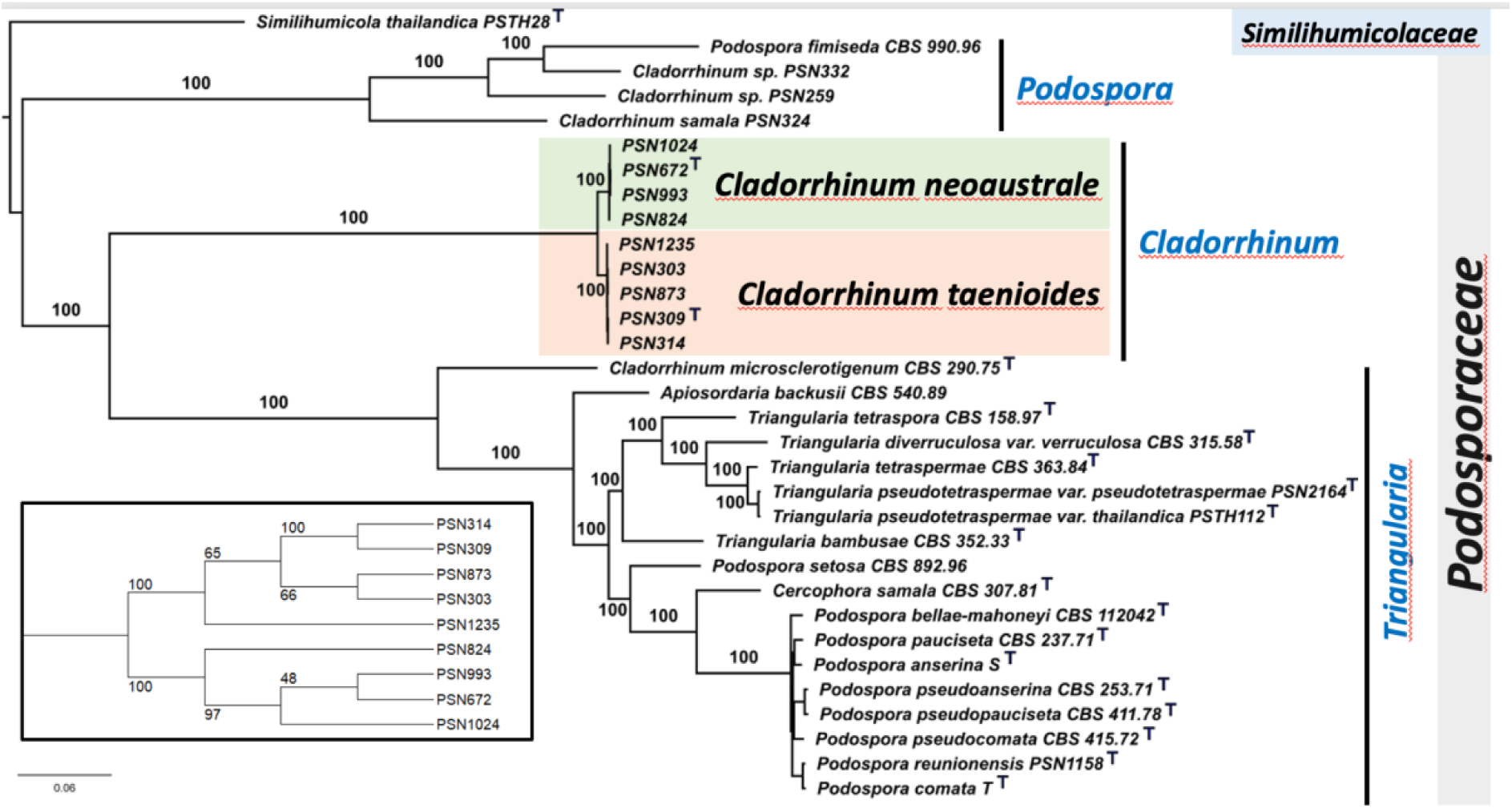
Phylogenomic tree of the *Podosporaceae* including *Cladorrhinum neoaustrale* and *C. taenioides* based on whole genome sequences. Thirty-two strains were included in the combined sequence analysis of 3,399 genes, which comprised 1,944,883 characters with gaps. The log-likelihood of the tree was -18236873.5326. The species names used are those of the genome sequence projects. The tree was rooted with *Similihumicola thailandica*, which belongs to the *Similihumicolaceae*. The three genera defined by (Wang et al. 2019) are indicated in blue. The ^T^ symbol indicates the type strains. Inset: cladogram of the *C. neoaustrale* and *C. taenioides* strains showing the bootstrap values.

Inter-sterility between the two *Cladorrhinum* species inferred here further supported their separation, by comparing interspecies and intraspecific crosses using two *C. taenioides* and two *C. neoaustrale* strains, using homokaryons of opposite mating types for each cross (Table S4). We found overall higher fertility in intraspecific crosses than in interspecies crosses. Indeed, perithecia were observed in all eight intraspecies crosses performed (within strains or between distinct strains), while perithecia were clearly observed at the frontier of parental thalli for a single one out of eight interspecies crosses performed (Chi2 test: d.f.=1; Pearson Chi2=14.32; P=0.0002). In addition, the single inter-specific cross with perithecia displayed lower perithecia density and nearly absence of spores compared to intra-specific crosses (Figure S2).

### A long-read-based genome assembly of *Cladorrhinum taenioides* PSN309

In order to investigate the presence of recombination suppression around the MAT locus, we generated a high-quality genome assembly of the wild *C. taenioides* PSN309 F0 MAT1-2 strain using long-read Oxford Nanopore sequencing. The long-read-based assembly was 40.6 Mb in size and contained seven contigs of more than 2.1 Mb each. Telomeric repeats were identified only at one end of two contigs (C_13 and C_14). The annotated transposable elements represented 7.7 % of the genome base pairs, with a majority of retrotransposons; for example, Gypsy LTRs (long-terminal repeat elements) represented 3.8% of the genome base pairs (Figure S3), as is also the case in *P. anserina* (Westerberg et al., 2024). Using the RIPper webtool analysis (Wyk et al. 2019) (ref), we detected low levels (<0.1%) of RIP signatures (repeat-induced point mutations), a fungal genome defense against transposable elements inducing C to T mutations in repeated sequences at target dinucleotides (Cambareri et al. 1989; Galagan and Selker 2004). The contig C_11, being 120 kb long with a low GC rate of 29.6%, is the mitochondrial genome.

### A standard MAT locus structure

The two *Cladorrhinum* species displayed standard ascomycete mating-type idiomorphs, separated in haploid genomes. We identified the MAT1-2 idiomorph by blast using the *P. anserina FPR1* (=*mat1-2-1*) gene as a query in the long-read-based assembly of *C. taenioides* PSN309 F0 MAT1-2. We found the MAT locus at 5.96 Mb on contig C_1, that is 9.49 Mb long and assembled from telomere to telomere. The MAT locus is located 1.72 Mb from the predicted centromere, whose position was inferred from high repeat density, low GC content and conserved location compared to the centromere of the *Podospora anserina* mating-type chromosome (at 4.24 Mb in contig C_1; Figure S4). We identified the MAT1-1 idiomorph by blast using the *P. anserina FMR1* (=*mat1-1-1*) gene as a query in the more fragmented assembly of *C. taenioides* PSN309 F0 MAT1-1 generated from Illumina reads: it was located on contig PSN309_1_1_C_350, that was only 27 kb long. The two mating-type idiomorphs span 3.7 kb and 4.6 kb, respectively, and display the canonical gene content of *Sordariales* MAT idiomorphs (Figure S5).

### No large-scale heterozygosity or rearrangement around the mating-type locus

We detected no large-scale recombination suppression in the reference *C. taenioides* genome PSN309, neither in terms of heterozygosity, rearrangements or transposable element content. To investigate footprints of recombination suppression around the MAT locus, we first assessed nucleotide sequence divergence between the MAT1-1 and MAT1-2 genomes of *C. taenioides* PSN309 F0 by examining the heterozygosity between the two genomes, particularly in the region surrounding the MAT locus. To this end, we called SNPs in *C. taenioides* PSN309 F0 MAT1-1 by mapping Illumina reads against the long-read-based assembly of *C. taenioides* PSN309 F0 MAT1-2. We detected 93 SNPs, which were located in a 169 kb region around the MAT locus, between 5,870 kb and 6,039 kb on contig C_1 (Figure 4). This region contained 48 predicted genes, with putative functions involved notably in mRNA translation, DNA replication, or several metabolic pathways (Table S5). We found only 6 SNPs in other genomic regions: 3 SNPs on the mating-type contig C_1, 1 SNP on contig C_2 and 2 SNPs on contig C_7. We detected no transposable element at all within the heterozygous region around the MAT locus. By comparing the *C. taenioides* PSN309 F0 MAT1-1 and MAT1-2 genome assemblies, we detected no chromosomal rearrangement in the region surrounding the MAT locus. However, the *C. taenioides* PSN309 F0 MAT1-1 genome, assembled from Illumina short reads, was highly fragmented, with four contigs identified in the heterozygous region flanking the MAT locus: contig PSN309_1_1_C_91 (121 kb), contig PSN309_1_1_C_350 (27 kb, carrying the MAT locus), contig PSN309_1_1_C_175 (65 kb), and contig PSN309_1_1_C_75 (121 kb). As a result, some rearrangements may have gone undetected.

**Figure 4.**
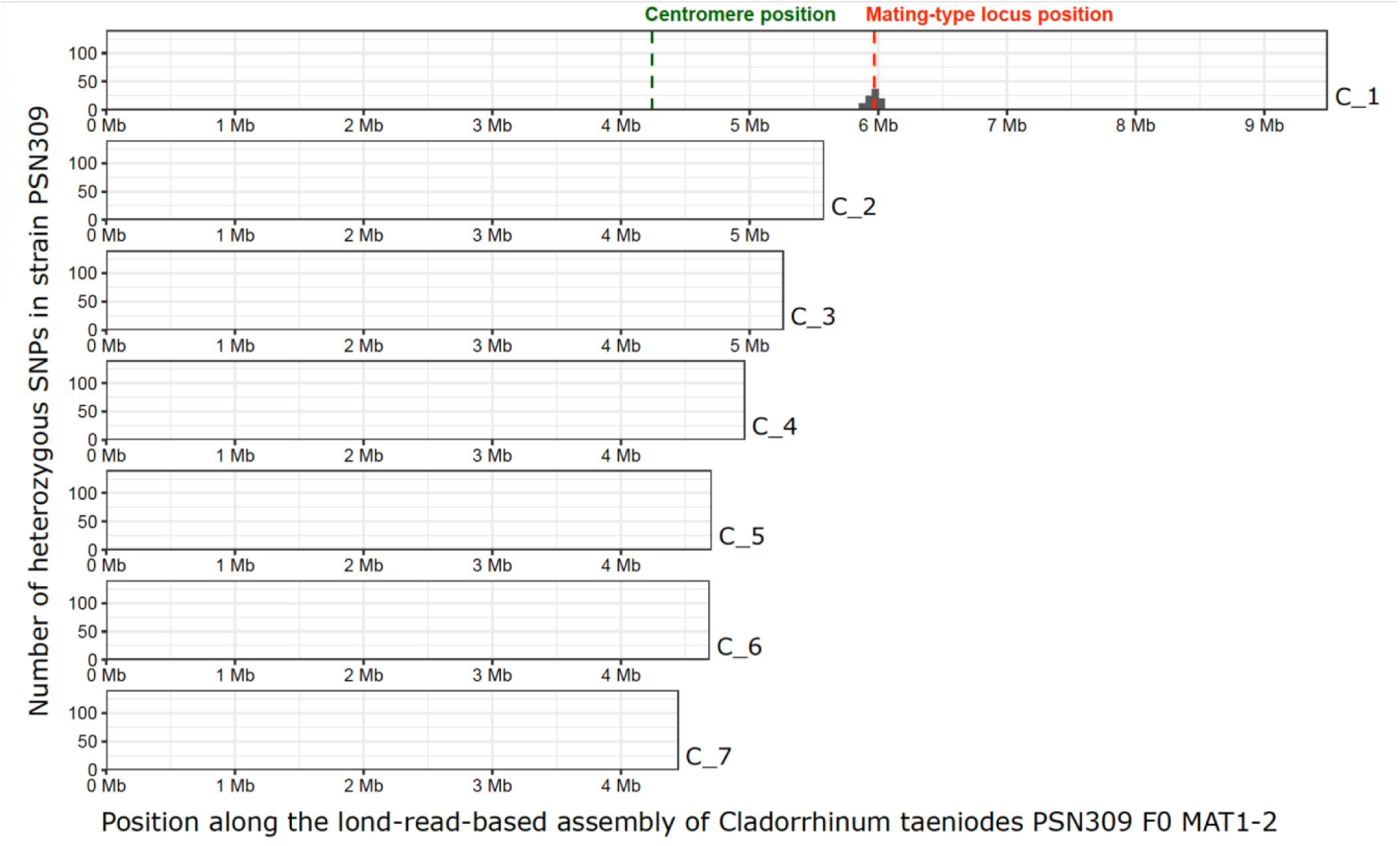
Genome-wide nucleotide divergence between the MAT1-1 and MAT1-2 genome of *Cladorrhinum taenioides* PSN309. The number of SNPs per 50-kb window between the MAT1-1 and MAT1-2 genomes of *C. taenioides* PSN309 was plotted along the MAT1-2 *C. taenioides* PSN309 long-read based genome assembly. Only the contigs larger than 1 Mb were represented. On contig C_1, the location of the mating-type (MAT) locus and the putative centromere are indicated respectively by a vertical dotted red line and vertical dotted green line, respectively.

Furthermore, we detected only small regions of heterozygosity around the MAT locus in other wild *Cladorrhinum* strains, except for one *C. neoaustrale* strain, which had a larger heterozygous region. In addition to the reference strain above, we investigated heterozygosity around the MAT locus in four other wild strains for which both MAT1-1 and MAT1-2 genomes were available: one additional *C. taenioides* strain (PSN1235) and three *C. neoaustrale* strains (PSN1024, PSN672, PSN824) (Figure S6). Heterozygosity was assessed by performing SNP calling using as a reference the long-read-based assembly of *C. taenioides* PSN309 F0 MAT1-2. In these strains, heterozygosity was mostly restricted to the region spanning 101–170 kb around the MAT locus (Figure S6; Table S6). Only *C. neoaustrale* PSN672 displayed a large heterozygous region, spanning 572 kb (Figure S6; 6; Table S6). Such large-scale heterozygosity found in a single strain could result from an outcrossing event a few generations ago, followed by a few generations of selfing, but with not enough selfing generations for homozygosity to be completely restored in recombining regions around the MAT locus.

Consistent with this hypothesis, the analysis of heterozygosity in the progeny of *C. neoaustrale* PSN672 after 10 generations of selfing revealed a substantial reduction of the heterozygous region around the MAT locus. By sequencing genomes of the mycelium resulting from 15 F11 heterokaryotic spores, we detected heterozygous regions ranging from 86 kb to 132 kb in length around the MAT locus. In all offspring, the extent of heterozygosity around the MAT locus was reduced relative to the F1 generation. A 86-kb interval remained consistently heterozygous in all F11 offspring, suggesting the presence of a small genomic region in which heterozygosity was maintained despite repeated selfing (Figure S7). This interval spans positions 5.928 Mb to 6.014Mb on contig 1 and encompasses the MAT locus. It contains at least a total of 130 SNPs differentiating the two mating-type haplotypes (Table S6).

### Tetrad analyses of *Cladorrhinum taenioides* PSN309 suggest a reduced diploid-like stage

By PCR genotyping the mating-type idiomorphs and by assessing the self-fertility of ascospore-derived mycelia, we inferred relatively low proportions of heterokaryotic ascospore-derived mycelia in *C. taenioides* PSN309 compared to previously studied pseudo-homothallic Sordariales (Silar 2020; Vittorelli et al. 2023). We studied a selfed cross of *C. taenioides* PSN309 (PSN309 F0 MAT1-2 × PSN309 F0 MAT1-1), one of the first strains that we had isolated in 2018, and from which we had isolated F1 tetrads (four-spore asci). We initially analysed one spore per tetrad using PCR amplification at the MAT locus on the 2 or 3-day-old ascospore-derived mycelium growing on petri dishes amended with G medium. Among the mycelia growing from the 74 analysed spores (referred to as PSN309 F1 progeny cross 1), 34 (46%) were heterokaryotic, while 40 (54%) were homokaryotic. No significant mating-type bias was observed among homokaryotic mycelia, with 18 (24.3%) identified as MAT1-1 and 22 (29.7%) as MAT1-2 (Pearson Chi²=0.40, d.f.=1, P=0.53).

To determine whether the relatively low proportion of heterokaryotic mycelia reflected the production of asci with four homokaryotic ascospores, due to the clustering of nuclei of carrying the same-mating, or subsequent changes in nuclear composition during the ascospores development or in growing mycelia from originally heterokaryotic ascospores, we analysed the three remaining spores from a subset of asci of the *C. taenioides* PSN309 F1 progeny. Indeed, if at least one spore was heterokaryotic, this means that all four spores in the ascus were heterokaryotic given mating-type segregation. Small spores (presumably haploid) were rarely observed, the vast majority of asci having four large spores. Ascospore-derived mycelia were again genotyped by PCR at the MAT locus (Figure 5A), and complete genotyping of the mycelia growing from all four ascospores was obtained for 26 tetrads. Across all ascospores of these fully characterised asci, 66 (63.5%) germinated into a heterokaryotic mycelium, 22 (21.1%) into a MAT1-1 homokaryotic mycelium, and 16 (15.4%) into a MAT1-2 homokaryotic mycelium. Only three asci yielded heterokaryotic mycelia from all their four spores, as assessed based on PCR-based genotyping of mycelia. For all analysed asci, however, at least one spore yielded a heterokaryotic mycelium (Figure 5A).

**Figure 5.**
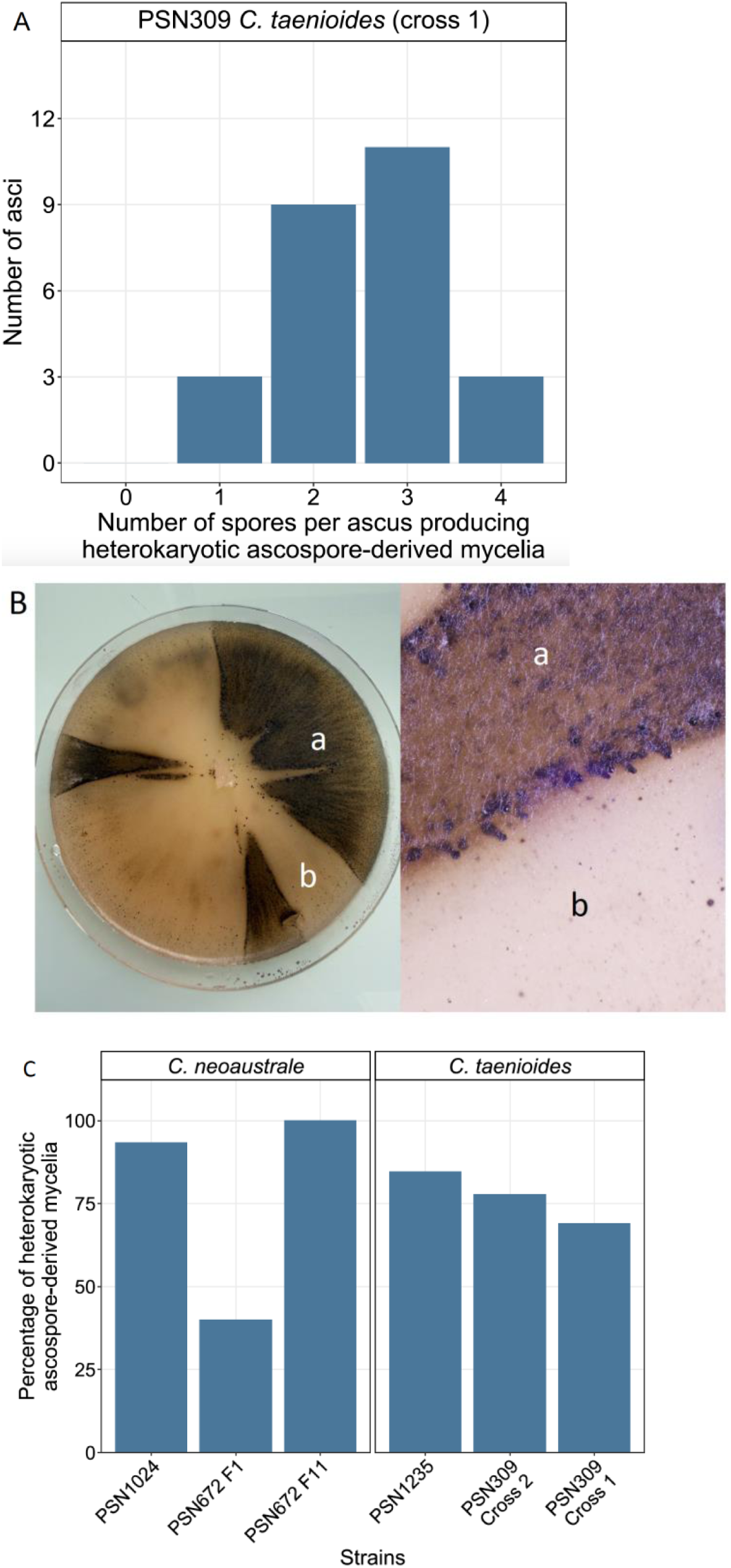
Percentage of heterokaryotic ascospore-derived mycelia in *Cladorrhinum taenioides* and *C. neoaustrale* strains. **A**. Number of spores per ascus that germinated to form heterokaryotic mycelia among 26 four-spored asci produced by crossing the *C. taenioides* PSN309 F0 MAT1-1 and PSN309 F0 MAT1-2 strains. MAT genotypes were assessed by PCR amplification of the MAT locus. **B.** Fungal culture of *C. taenioides* PSN309 on a Petri dish containing oat agar (OA) medium, showing sectors of fertile mycelium (a) and sterile mycelium (b). The right part is a zoom at a frontier between fertile (a) and sterile (b) sectors. **C.** Results of the progeny analysis of the *C. taenioides* PSN309 and PSN1235 strains and the *C. neoaustrale* PSN1024 and PSN672 strains. For each cross, the percentage of heterokaryotic ascospore-derived mycelia is shown, as assessed by self-fertility tests. For the PSN309 strain, two different crosses were studied. For the PSN672 strain, both F1 and F11 selfed progeny were studied.

**Figure 6.**
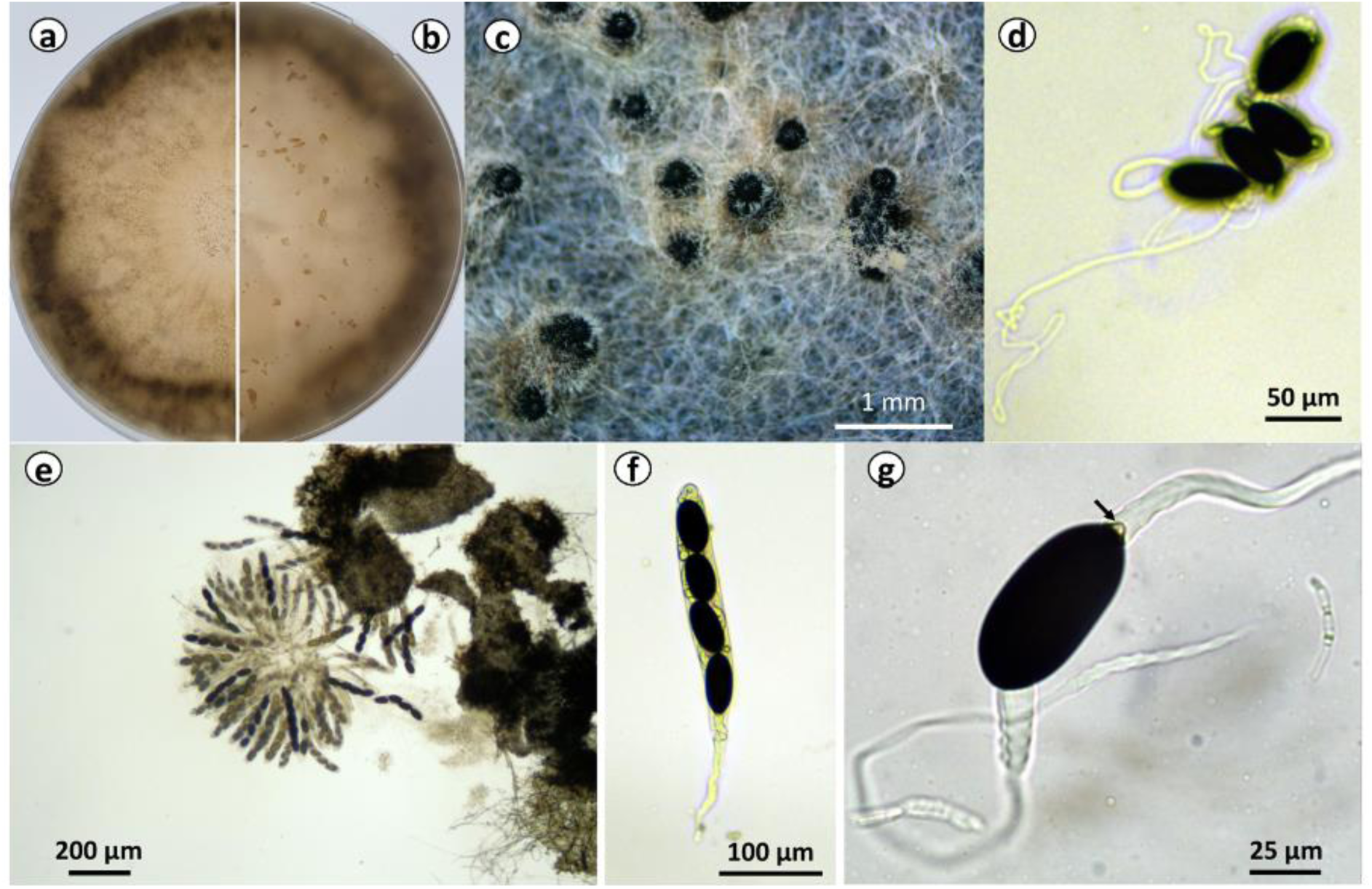
Morphology of *Cladorrinum neoaustrale* (Speg.) Silar nom. nov. strain PSN672. Mycelium on oat agar (OA) medium, top view (a) and bottom view (b). Perithecia on OA (c). Ascospores expelled on water agar showing the long secondary appendages (d). Rosettes of asci (e). Ascus (f). Ascospore with the arrow showing the primary appendage (g).

**Figure 7.**
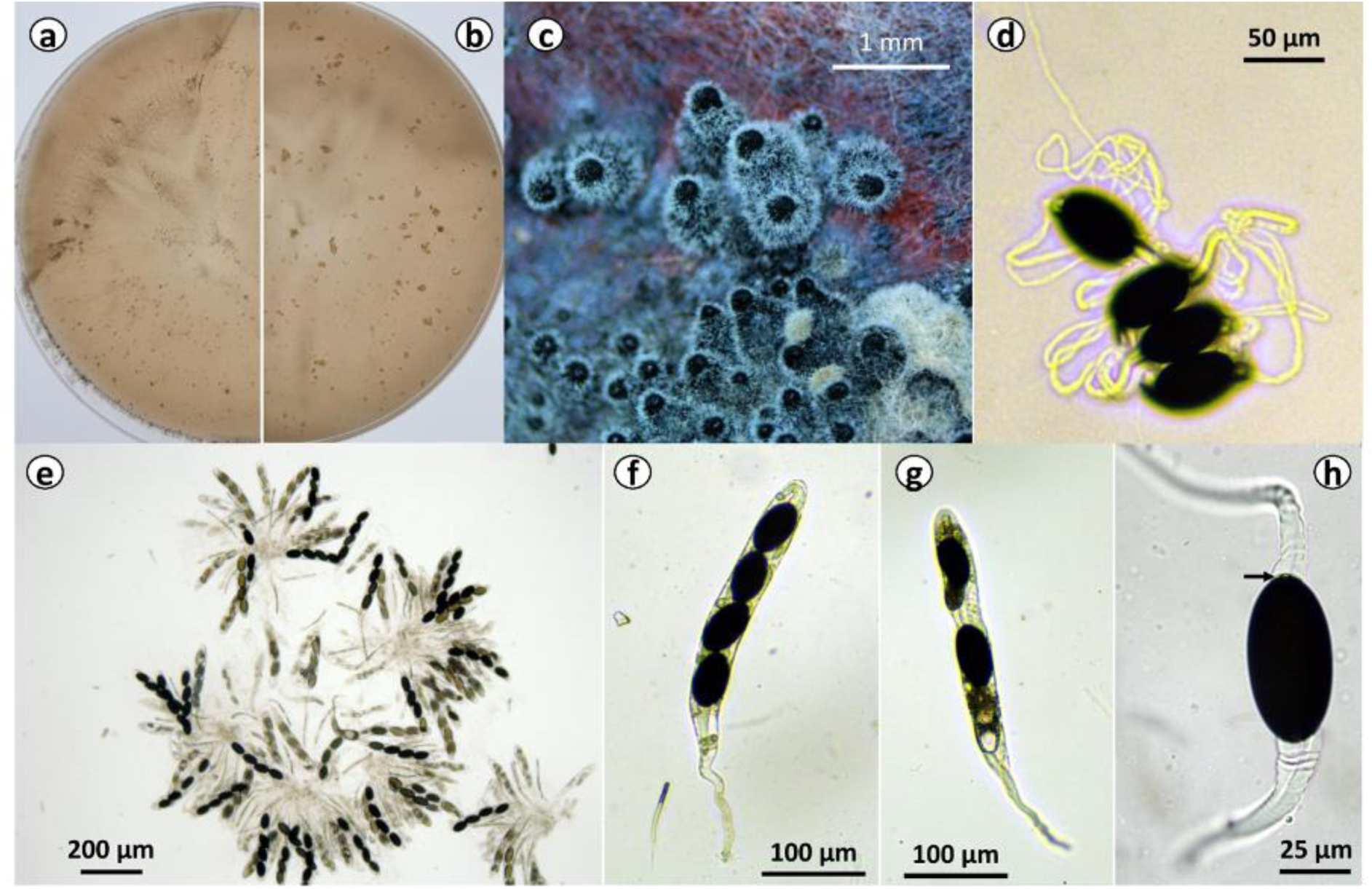
Morphology of *Cladorrinum taenioides* (Griff.) Silar comb. nov. strain PSN309. Mycelium on oat agar (OA) medium, top view (a) and bottom view (b). Perithecia on OA (c). Ascospores expelled on water agar showing the long secondary appendages (d). Rosettes of asci (e). Normal ascus (f). Abnormal ascus (g). Ascospore with the arrow showing the minute primary appendage (h).

These observations are consistent with the formation of four heterokaryotic spores following meiosis, followed by substantial losses of nuclei of one mating type during spore formation or subsequent mycelial growth (Fig. S1B, scenarios 1–2), as previously observed when genotyping four-spore asci in the *S. tetrasporum* species complex (Vittorelli et al. 2023; Guyot et al. 2025). Alternatively, some mycelia scored as homokaryotic by PCR may have retained nuclei of both mating types, but with one mating type strongly under-represented among the nuclei, and therefore below the PCR detection threshold (Fig. S1B, scenario 3).

The hypothesis that nuclear composition can change during vegetative growth is also consistent with our observations of sectoring in the *C. taenioides* PSN309 selfed progeny (Figure 5B), as observed previously in the *S. tetrasporum* species complex (Guyot et al. 2025). Growing colonies frequently developed sectors with distinct morphologies, and only one sector type produced perithecia. This pattern is consistent with the loss of nuclei of one mating type in sterile sectors, whereas sectors retaining both mating types remain self-fertile and perithecia can be observed at the border of the fertile and sterile sectors. Supporting this interpretation, we observed the same distinct mycelial morphologies between homokaryons of opposite mating types in *C. taenioides* PSN309 (Fig S7A). Such losses of nuclei of one mating type may also explain the single MAT idiomorph detected in genomes for wild strains for which the initial cultures produced perithecia, indicating that were therefore heterokaryotic: the subsequent sub-cultures for DNA extraction have likely lost nuclei of one mating type. Alternatively, as noted above, nuclei of one mating type may have become sufficiently rare to escape PCR detection without being completely lost (Fig. S1B, scenario 3).

We therefore used self-fertility as an additional way to assess heterokaryosis, looking for perithecia production in standardized Petri dish assays. Among mycelia growing from 68 ascospores previously genotyped with PCR from the *C. taenioides* PSN309 F1 progeny, 47 were self-fertile (69%; Figure 5C, cross 1), including 21 that were not detected as heterokaryotic by PCR. This discrepancy suggests that either PCR genotyping can underestimate the proportion of heterokaryotic mycelia or that sub-culturing steps of the same mycelium led to different results for the same F1 progeny due to nucleus loss during growth of some cultures and not others. Nevertheless, 31% of the tested mycelia were not self-fertile in the standardized Petri dish assay. Together with the observed sectoring, this result is consistent with substantial loss of nuclei of one mating type during mycelial growth.

### Variable percentages of heterokaryotic ascospore-derived mycelia among further *Cladorrhinum taenioides and C. neoaustrale* strains

After having isolated in 2021 further *Cladorrhinum* strains that produced perithecia in the lab, we also investigated the extent of heterokaryosis in their mycelia, based on self-fertility assays that can be easily applied to multiple strains. We also re-included *C. taenioides* PSN309 in this assay to compare results across strains in the same experiment. We found substantial variation among the four strains studied and the F11 progeny described above (Table S7). Among the mycelia grown from ascospores, a very high proportion was self-fertile in the selfed progeny of *C. neoaustrale* PSN1024 (97%; Figure 5C), while lower, yet still substantial, proportions were self-fertile in other strains: 78% in the second cross of *C. taenioides* PSN309 F0 MAT1-2 and PSN309 F0 MAT1-1 (PSN309 F1 cross 2), 85% in the selfed progeny of *C. taenioides* PSN1235, and 40% in the selfed progeny of *C. neoaustrale* PSN672 (Figure 5C). In this selfed progeny of *C. neoaustrale* PSN672, however, we also observed the presence of different sectors, with morphological differences (Figure S7B (1)). No sectors with distinct morphologies were observed in the mycelium of the selfed progeny of *C. neoaustrale* PSN1024 and *C. taenioides* PSN1235 (Figure S7). Conversely, in the F11 progeny of *C. neoaustrale* PSN672, the percentage of tested ascospore-derived mycelia that were self-fertile increased to 100% and no sectors were observed.

### Taxonomy

Based on the ANI values (Table 2) and the phylogenetic trees (Figures 2 and 3), two species belonging to the *Cladorrhinum* genus in the *Podosporaceae* family need to be formally defined. We thus introduce two new combinations *C. neoaustrale* and *C. taenioides*. There is already a species named *C. australe* (IF 813037), making it necessary to change the epithet for *C. neoaustrale.* Because the types for *P. taenioides* and *P. australis* are of historical value and cannot be used for molecular identification, we epitify the two species by providing lectotypes, epitypes and ex-type cultures.

***Cladorrhinum neoaustrale*** (Speg.) P. Silar **nom. nov.** IF= 905861 (Figure 4)

**Basionym: *Hypocopra australis*** Speg., Anal. Soc. Cient. Argent. 10(3): 137 (1880) IF= 150020

**Replaced synonym: *Podospora australis*** (Speg.) Niessl, Hedwigia 22: 156 (1883) IF= 100929.

***Lectotypification*** (Index Fungorum IF= 150020): Buenos Aires, on dung of *Myopotamus coypa, Lectotypus hic designatus*.

***Epitypification*** (Index Fungorum IF= under request): FRANCE, Camargue. Isolated from horse dung in 2021. (PC0820509 *Epitypus hic designatus*); ex-epitype culture: strain PSN672 deposited at the CIRM-CF culture collection (https://www.cirm-fungi.fr/) # BRFM 3916, Figure 4. The whole genome sequence of PSN672 is available in GenBank under BioProject under request.

Materials examined: strain PSN672 isolated from horse dung collected in Camargue, France in 2021, strain PSN824 isolated from Bertre in the Tarn department, France in 2021 from hare dung, strain PSN1024 isolated from horse dung collected in St Remy de Provence, France in 2021. Genome sequences for the three strains are available under the BioProject under request in GenBank.

Sexual morph: ostiolate perithecia diam. 600 ± 100 µm (n=10), covered with numerous short straight hairs including the base of the neck; peridium membranaceous or slightly coriaceous, dark brown and opaque, *textura* not distinctly visible. Asci 4-spored, cylindrical with a very long stipe. Ascospores uniseriate at first unicellular then bicellular with the spore head directed towards the apex of the ascus. Spore head elongate-ellipsoid, 48.4 ± 2.4 × 24.3 ± 1.5 µm (n=50); primary appendage reduced to very small basal apiculum with a semi-discoidal shape, clearly visible in most ascospores, 3.7 ± 0.4 × 2.2 ± 0.6 (n=8) µm; very long secondary taenioid appendages at both poles of the spore head, with a width of 8.8 ± 0.8 µm (n=10) at the junction with the ascospores; the apical one is slightly off-center while the other is centered around the primary appendage covering it completely. Asexual morph: *Cladorrhinum*-like likely producing spermatia rather than conidia. Cultural characteristics: mycelium growing at 3.3 mm/day on M2, 4.0 mm/day on OA, V8 and carrot juice agar. Perithecia were differentiated in large numbers on OA, in fewer numbers on M2 and M0 + miscanthus, and none on carrot juice agar and V8. Ascospores germinated efficiently on germination (G) medium supplemented with 5 g/L of yeast extract treated with a heat shock at 63°C for 30 minutes.

Notes – Morphology of PSN672, PSN824 and PSN1024 is very similar (for PSN824: perithecium diam. 530 ± 70 µm, ascospore head 52.2 ± 2.8 × 25.7 ± 1.0 µm, primary appendage 3.2 ± 0.5 × 1.2 ± 0.3 µm, secondary appendage 9.6 ± 0.9 µm, for PSN1024: perithecium diam. 670 ± 90 µm, ascospore head 49.7 ± 2.3 × 25.5 ± 1.7 µm, primary appendage 3.7 ± 0.3 × 2.1 ± 0.4 µm, secondary appendage 9.6 ± 0.7 µm) and is consistent with the original description of *Hypocopra australis* by C. Spegazzini (Spegazzini 1880), although the ascospores appear slightly shorter in our isolates (Spegazzini ascospores are 60-70 x 25-30 µm). Note that the protologue did not describe the primary appendage. Morphology is also consistent with the drawing of *P. australis* by (Mirza and Cain 1969); in particular, the shapes and sizes of the primary appendages of PSN672 and PSN1024 are identical to the description given by Mirza and Cain (4 × 1-5 µm), while the ones of PSN824 are slightly smaller; however our ascospores are bigger (Mirza & Cain give 26 -34 x 12-20 µm; (Mirza and Cain 1969)). They are nonetheless in the size range given by N. Lundqvist for the species (Lundqvist 1972): (46-)50-65 × (25-)27-36(-37) µm. The drawing of the species by N. lundqvist also shows a conspicuous primary appendage of 2.5-3 µm. Phylogenomic analyses place this species close to *C. taenioides* and *Cladorrhinum foecundissimum*, the type species of the genus *Cladorrhinum*. Mirza and Cain considered *P. australis* and *P. taenioides* as synonyms (Mirza and Cain 1969); the two species are reported as such in the Mycobank database, but considered as separate in the Index Fungorum and Fungal Names databases. We segregate *C. neoaustrale* and *C. taenioides* into different species on the basis of their differentiated genome sequences. Additionally, *C. neoaustrale* is much more fertile on M2 and M0+miscanthus than *C. taenioides*. We also provide epitypification of the species due to the lack of a holotype usable for molecular identification.

***Cladorrhinum taenioides*** (Griffiths) P. Silar **comb. nov.** IF= 905834(Figure 5)

***Lectotypification*** (Index Fungorum IF= 228455): On dung of *Equus caballus* and of *Bos taurus*: New Jersey, New York, New Mexico, Montana, Arizona, Texas, South Dakota, Mississippi, Alabama, Rhode Island Griffiths, Aug. 1899; Griffiths, Sept. 1899; Banker, Aug. 1899; Wooton, Jan. 1900; Demeterio, Aug. 1899; Tyler, Jan. 1900; Long, Jan. 1900; Carter, Nov. 1899; Tracy, Sept. 1899; Earle, Aug. 1899; Underwood, Nov. 1899; Griffiths, Sept. 1900*, Lectotypus hic designatus*.

***Epitypification*** (Index Fungorum IF= under request): ITALY, Cassimorenga, Apennines. Isolated from rabbit dung in 2018*. (*PC0820511*, epitypus hic designatus*); ex-epitype culture: strain PSN309 deposited at the CIRM-CF culture collection (https://www.cirm-fungi.fr/) n° BRFM 3919, Figure 5.

The whole genome sequence of PSN309 is available in GenBank under BioProject under request.

**Materials examined:** strain PSN309 isolated from rabbit dung collected in Cassimorenga, Appennines, Italy in 2018, and strain PSN1235 isolated from cow dung collected in Montpellier, France in 2022. Genome sequences for the two strains are available under BioProject under request in GenBank.

Sexual morph: ostiolate perithecia diam. 550 ± 60 µm (n=10), covered with numerous short straight hairs including the base of the neck; peridium membranaceous or slightly coriaceous, dark brown and opaque, *textura* not distinctly visible. Asci 4-spored, cylindrical with a very long stipe. Ascospores uniseriate at first unicellular then bicellular with the spore head directed towards the apex of the ascus. Spore head elongate-ellipsoid, 53.9 ± 4.1 × 28.8 ± 2.3 µm (n=50); primary appendage reduced to a tiny basal apiculum with a semi-discoidal shape, sometime invisible, 2.9 ± 0.3 × 0.9 ± 0.3 (n=5 + 3 ascospores without a discernible primary appendage) µm; very long secondary taenioid appendages at both poles of the spore head, with a width of 11.5 ± 1.6 µm (n=10) at the junction with the ascospores; the apical one is slightly off-center while the other is centered around the primary appendage covering it completely. Asexual morph: *Cladorrhinum*-like likely producing spermatia rather than conidia. Cultural characteristics: mycelium growing at 3.3 mm/day on M2, 4.4 mm/day on OA, V8 and carrot juice agar. Perithecia were differentiated in large numbers on OA, M2 and M0 + miscanthus, in fewer numbers on carrot juice agar and none on V8. Ascospores germinated efficiently on the germination (G) medium supplemented with 5 g/L of yeast extract treated with a heat shock at 63°C for 30 minutes.

Notes – The morphologies of PSN309 and PSN1235 were very similar (for PSN1235: perithecium diam. 650 ± 70 µm, ascospore head 56.4 ± 5.4 × 30.6 ± 4.8 µm, primary appendage 3.5 ± 0.2 × 1.1 ± 0.2 µm, not visible in some ascospores, secondary appendage 11.1 ± 0.9 µm). This is consistent with the original description of *Pleurage taenioides* by D. Griffiths (Mirza and Cain 1969), his drawing of the species being strikingly similar to our observations, especially the minute size of the primary appendage that is barely visible in the two strains that we examined. PSN309 presents a fair number of abnormal asci with varying numbers of aborted ascospores (Fig. 5), which were not seen in PSN1235. The reason for this atypical feature is not known, but it may be due to the presence of genetic incompatibilities or spore killers. Phylogenomic analyses place this species close to *C. neoaustrale* and *Cladorrhinum foecundissimum*, the type species of the genus *Cladorrhinum*. Mirza and Cain considered *P. taenioides* and *P. australis* as synonyms; the two species are reported as such in the Mycobank database, but considered as separate in the Index Fungorum and Fungal Names databases. We segregate *C. taenioides* and *C. neoaustrale* into different species on the basis of their differentiated genome sequences. Moreover, *C. neoaustrale* is much more fertile on M2 and M0+miscanthus than *C. taenioides*. We also provide epitypification of the species due to the lack of a holotype usable for molecular identification.

## Discussion

The main objective of this study was to contribute testing the association between an extended dikaryotic life stage and the suppression of recombination around the MAT locus, using the formerly known *P. australis* as an additional pseudo-homothallic lineage, potentially representing an independent evolution of pseudo-homothallism relative to previously studied systems (Jay et al. 2024; De Filippo et al. 2026). According to some recent theoretical models, the evolution of recombination suppression on sex chromosomes and mating-type chromosomes is explained by effects emerging from the presence of recessive deleterious mutations in genomes (Jay et al. 2024; Jay et al. 2025; Guyot et al. 2026). Recombination suppression should then only evolve in lineages with a predominant diploid-like life-cycle. Recombination suppression around mating-type loci would in particular be predicted to evolve only in pseudo-homothallic (diploid-like) Sordariales fungi, as heterothallic lineages are haploid (or homokaryotic) during most of their life cycle. These models, however, do not predict an evolution of recombination suppression in all diploid-like life cycles, as it also depends on the importance of the diploid-like phase and on other parameters, such as the mutation rate and the recombination rate (Jay et al. 2024; Jay et al. 2025; Guyot et al. 2026). To contribute testing the hypothesis of an association between pseudo-homothallism and recombination suppression in the MAT-locus flanking regions, we assessed the importance of the diploid-like phase in the lineage known as *P. australis,* considered pseudo-homothallic, and tested whether it displayed recombination suppression around its mating-type locus; for this goal, we used genomic analyses and studies of the self-fertility of ascospore-derived mycelia.

### Cryptic species

While *P. australis* and *P. taenioides* are considered synonyms in the Mycobank database, they are considered as distinct species in the Index Fungorum and Fungal Names ones (accessed on June 9th 2026). Mirza and Cain state that they “have seen the type material of this species [*P. australis*] and found it to be the same as *P. taenioides”* (Mirza and Cain 1969). They also state that a third species, described as *Sordaria macrura* Bayer, is also a synonym. Lundqvist concurs that the three species are synonyms, but states that the holotype for *S. macrura* has been lost and that only one slide of all *P. taenioides* specimens remains. He concludes that this slide “becomes the natural lectotype for *P. taenioides*” (Lundqvist 1972). These holotype losses make it difficult to ascertain which species were originally described. Moreover, several authors noted that the species is highly variable in ascospore size and shape, the primary appendage development and the presence of hair on the perithecium (Mirza and Cain 1969; Lundqvist 1972; Doveri 2004). Lundqvist however states that its “Swedish material is homogeneous”.

In our study, as typically the case in fungi, genomic analyses revealed the existence of cryptic species within the morphological species *P. australis,* and we described two species, that we assigned to the *Cladorrhinum* genus based on their position in phylogenetic trees. Our results indeed provide convergent evidence supporting the recognition of *Cladorrhinum neoaustrale* and *C. taenioides* as distinct species rather than synonyms. Although the two taxa exhibited only limited divergence at the ITS locus, whole-genome comparisons revealed two well-separated genetic groups, with high intraspecific ANI values (>99.8%) and markedly lower interspecific similarities (97.7–97.8%), a pattern further supported by both ITS+LSU and phylogenomic analyses. The two species differed in their growth and reproductive characteristics in culture, with *C. neoaustrale* displaying substantially higher fertility on M2 and M0 medium supplemented with miscanthus. Furthermore, interspecific crosses were overall much less fertile than intraspecific ones.

All authors having previously studied *P. australis* or *P. taenioides* indicated that the species were cosmopolitan (Mirza and Cain 1969; Lundqvist 1972; Doveri 2004). The strains isolated here came from the South of France and the Apennines in Italy. We found so far a single species in Italy (*C. taenioides*), and both in the south of France, showing an overlap in their geographical distribution. In the dozens of dung samples originating from northern France analyzed previously (Silar et al. 2025), neither of these two species were found. This could suggest that both species identified here prefer warmer climates, but dried specimens considered to belong to *P. australis* in collections came from northern and eastern Europe and Canada (Lundqvist 1972), regions that are colder than southern France. All of our strains were isolated from herbivore dung, and none from soil, although numerous soil samples were also screened for Sordariales (Silar et al. 2025), indicating a narrow ecological niche for these two species. Together, these differences in growth and reproductive characteristics in culture indicate that the taxa previously grouped under the names *P. australis* and *P. taenioides* likely represent a species complex comprising several independently evolving lineages.

Our phylogenomic analyses show that the *Cladorrhinum* genus branches with 100% confidence as sister group to the *Triangularia* genus defined by (Wang et al. 2019), with *Podospora* as defined by Wang et al. being the closest outgroup, confirming the position for the genus determined with low confidence by their DNA barcodes. The taxonomy of the *Podosporaceae* is currently unclear. Wang et al. recently split the family into three genera: *Podospora*, the type genus, *Triangularia* and *Cladorrhinum*. This resulted in many species changing names, while others awaiting taxonomic revisions and still carrying their old names. For example, the genus *Cladorrhinum* is now well defined thanks to molecular data, but many genome projects still use the old names for some “*Cladorrhinum”* strains that do not belong to the newly defined *Cladorrhinum* genus. Moreover, the taxonomic revision by (Wang et al. 2019) resulted in the renaming of the only well-studied species of the family, from *P. anserina* to *T. anserin*a, a change that was met with strong resistance from the research community working on this model organism (Ament-Velásquez et al. 2020; Silar 2020; Bhunjun et al. 2024). Previous analyses showed that the DNA barcodes used for phylogenetic analyses were not sufficient to accurately define the generic relationships within the family, and *Podospora* and *Triangularia* were synonymized (Ament-Velásquez et al. 2020). Additionally, it has been proposed to change the type of the *Podospora* genus from *P. fimiseda* to *P. anserina* (Vogan et al. 2021). Pending the decision on the type change, the taxonomy of the family remains unresolved.

The two species described here displayed similar life cycles, both exhibiting some features of pseudo-homothallism, but with the vegetative growth phase not always remaining diploid-like. In both taxa, small spores were rarely observed and the overwhelming majority of asci had four spores, suggesting that haploid spores are uncommon. However, nucleus loss appears to be frequent during mycelium growth, regularly leading to homokaryotic mycelia, as observed in *Schizothecium* fungi (Vittorelli et al. 2023; Guyot et al. 2025). The frequent appearance of sectors with contrasting mycelium morphologies might be due to some physiological differences between the homokaryotic and heterokaryotic mycelia, or to the presence of genetic interactions between the two nuclei in the heterokaryons, even if there were few genetic differences between *MAT1-1* and *MAT1-2* nuclei within strains. In particular, some recessive deleterious mutations may be present in the small heterozygous region around the MAT locus, as shown to be present in other Sordariales fungi (Guyot et al. 2025), and would be sheltered in heterokaryons but not in homokaryons.

### Absence of large-scale recombination suppression

Our genomic analyses indicated an absence of large-scale recombination suppression in regions flanking the MAT locus in the two *Cladorrhinum* species studied here. Indeed, we only detected a relatively small region of nucleotide divergence between mating types (∼97 - 170 kb), which may reflect either a localized reduction in recombination or residual heterozygosity generated by outcrossing events some generations ago, followed by several generations of selfing. Indeed, sites close to the mating-type locus are expected to return to homozygosity more slowly than the rest of the genome under selfing regimes (Hartmann et al. 2021ab). The near absence of polymorphism across the rest of the genome is consistent with a recent history dominated by selfing in the *Cladorrhinum* fungi studied here, in agreement with pseudo-homothallism. The large region of heterozygosity around the MAT locus in *C. neoaustrale* PSN672 was substantially reduced after 10 generations of experimental selfing events. This reduction supports the absence of large-scale recombination suppression, as it indicates that recombination occurs in the region. It also suggests that part of the heterozygosity resulted from former outcrossing events, consistent with occasional outcrossing in nature. Moreover, we found no evidence of chromosomal rearrangements or transposable element enrichment near the mating-type locus. The putative non-recombining region in the two *Cladorrhinum* fungi, if any, is thus markedly smaller (<200 kb) than the extensive regions reported in other pseudo-homothallic ascomycetes (>600 kb), such as *N. tetrasperma* (Menkis et al. 2008; Sun et al. 2017), *P. anserina* (Grognet et al. 2014; Hartmann et al. 2021a; Silar et al. 2025), and *Schizothecium* fungi (Vittorelli et al. 2023; De Filippo et al. 2026).

### Apparently lower frequencies of ascospore-derived mycelia compared to other pseudo-homothallic species

Genotyping of the MAT locus and self-fertility tests in ascospore-derived mycelia supported the existence of an extended heterokaryotic phase in the life cycle. The observed proportion of heterokaryotic mycelia (as low as 40%, Table S7) appeared however lower than values reported in other pseudo-homothallic ascomycetes, e.g. 87% of heterokaryotic mycelia germinating from ascospores in *S. tetrasporum*; (Vittorelli et al. 2023); 99% in *P. anserina* (Silar 2020). We also found variation between strains and even between a strain and its F11 progeny.

The existence of homokaryotic ascospore-derived mycelia may in principle be due to the production of homokaryotic spores associated with particular mating-type segregation or to the loss of nuclei of one mating type during spore development or mycelium growth (Fig S1). The segregation of the mating-type locus at meiosis and the cytological pattern of spore formation remain to be investigated in these pseudo-homothallic *Cladorrhinum* fungi, in particular whether frequent second-division segregation occurs at meiosis for the mating-type locus (Fig S1A), as found in other pseudo-homothallic fungi (Marcou et al. 1979; Vittorelli et al. 2023). The presence of at least one heterokaryotic spore in tetrad analyses, the observation of sectors with different morphologies and self-fertility abilities, and the detection of a single mating type in ascospore-derived mycelia, altogether suggest that ascospores are initially heterokaryotic and that losses of nuclei of one mating type regularly occurs during mycelium growth, as previously described in *S. tetrasporum* (Vittorelli et al. 2023; Guyot et al. 2025). The size of the ascospores in these *Cladorrhinum* fungi was larger (about 55×30 μm for the spore head) compared to previously studied pseudo-homothallic fungi (about 35×25 μm in *P. anserina* (Silar 2020) and about 25×15 μm in *Schizothecium* fungi (De Filippo et al. 2025)). This may correspond to a higher number of mitoses during spore-head formation. Indeed, cell volume is often correlated with the number of nuclei that they contain. This could thus open the possibility of a stronger bias in the amplification of nuclei of one particular mating type during ascospore maturation (Fig. S1B). Differences in nucleus ratios of the two mating types have also been documented in *Neurospora tetrasperma* (Samils et al. 2014).

For a pseudo-homothallic life cycle to allow reproductive insurance by selfing, not only should the ascospores pack two nuclei carrying compatible mating types, but the mycelium should stay heterokaryotic until sexual reproduction. A possibility is that the nucleus loss observed here is a kind of laboratory artefact, sexual reproduction and death occurring after a shorter time of mycelial growth in nature, before any nucleus loss; this may be the case in the dung where these fungi live, being ephemeral and small habitats. Alternatively, occasional nucleus loss could be beneficial in nature, by allowing some outcrossing events, which would correspond to a genuine partial pseudo-homothallism; this would be consistent with the long tract of heterozygosity found in the strain *C. neoaustrale* PSN672, that was reduced after 10 generations of selfing; all other strains studied were, however, highly homozygous genome-wide. It may also be that the species studied here have recently evolved pseudo-homothallism, and not optimized the maintenance of two nuclei during mycelium growth. If their life cycle is in fact only partially diploid-like in nature, this could weaken selection for recombination suppression across large regions, as predicted under the models involving recessive deleterious mutations, because even the recessive deleterious alleles closely linked to the MAT locus would not remain sheltered throughout the entire lifecycle. Such a mixed reproductive system, with both self-fertile and self-sterile mycelia, may represent a compromise between the assurance of successful sexual reproduction in ephemeral dung habitats and the generation of genetic variation by outcrossing.

## Conclusion

We describe here two cryptic species within the morphological species previously known as *P. australis;* we reassigned them to the genus *Cladorrhinum*, and we confirmed that they exhibit several features of pseudo-homothallic ascomycetes. However, we observed patterns that seem to differ from other pseudo-homothallic Sordariales fungi. Indeed, the *Cladorrhinum* strains studied appeared to display lower proportions of heterokaryotic ascospore-derived mycelia than in other pseudo-homothallic Sordariales fungi. In addition, we did not detect any large-scale recombination suppression around the MAT locus in these lineages. These findings are consistent with the recent theoretical models based on the effects of recessive deleterious mutations in genomes predicting that recombination suppression around a permanently heterozygous locus can only evolve in species with a life cycle being predominantly diploid-like (Charlesworth and Wall 1999; Antonovics and Abrams 2004; Jay et al. 2024; Jay et al. 2025; Guyot et al. 2026). Indeed, in the *Cladorrhinum* species studied here, the heterokaryotic phase may not be sufficiently predominant across strains and over time for recombination suppression to evolve. The time spent as homokaryotic mycelium may indeed be too high, allowing the purging of recessive deleterious mutations and preventing their sheltering. However, observations were variable across strains and generations and it remains to be assessed how frequent are heterokaryons in nature. Alternatively, the absence of large-scale recombination suppression may be due to other species-specific important features that are not conducive to the evolution of recombination suppression. Indeed, the theoretical models based on the effects of recessive deleterious mutations in genomes also highlight the importance of other parameters, such as the mutation rate and the recombination rate (Charlesworth and Wall 1999; Antonovics and Abrams 2004; Jay et al. 2024; Jay et al. 2025; Guyot et al. 2026). It may also be that pseudo-homothallism evolved too recently in this group, with thus not enough time for recombination suppression to evolve. Importantly, our results also decouple the evolution of pseudo-homothallism from that of recombination suppression, showing that there is no fundamental developmental constraint on their joint evolution.

## Material and Methods

### Strain collection and isolation

Growth conditions, media and genetic methods were those used for the model fungus *Podospora anserina* (Rabenh.) Niessl (Silar 2020). M0 minimal base medium has the following composition: KH2PO4 0.25 g/L, K2HPO4 0.3 g/L, MgSO4/7H2O 0.25 g/L, Urea 0.5 g/L, Thiamine 0.05 mg/L, Biotine 0.25 μg/L, Citric Acid 2.5 mg/L, ZnSO4 2.5 mg/L, CuSO4 0.5 mg/L, MnSO4 125 μg/L, Boric Acid 25 μg/L, Sodium Molybdate 25 μg/L, Iron Alum 25 μg/L, Agar 12 g/L. It is supplemented with potato dextrin at 4 g/L to yield the M2 minimal medium or with 0.5 g/plate of shredded miscanthus to yield M0+ Miscanthus. The OA medium contains 20 g/L of Quaker oat meal and 12 g/L of agar. The V8 and carrot juice media were made by mixing and autoclaving 200 mL of V8 juice or carrot juice, respectively, with 2 g of CaCO3, 15 g of agar and 800 mL of water. G germination medium is ammonium acetate 4.4 g/L, bactopeptone 15 g/L, and agar 12 g/L. G + yeast extract medium is G medium amended with yeast extract 5 g/L and agar 12 g/L. From all tested media and conditions, incubation on OA at 27°C with constant illumination was found to be the best condition for perithecium production. Germination effectiveness was best achieved on the germination medium supplemented with 5 g/L of yeast extract, with a heat shock of 30 minutes at 63°C before an overnight incubation at 27°C.

The PSN303, PSN309 and PSN314 strains were recovered from germinating whole four-spored asci. For PSN672, PSN824, PSN873, PSN993, PSN1024 and PSN1235, the four individual ascospores originating from a single ascus were germinated separately. For these later strains, the four isolates were labelled PSNxxxA, PSNxxxB, PSNxxxC, PSNxxxD. All asci were expelled from perithecia fruiting on dung and ascospores were transferred to germination medium supplemented with 5 g/L of yeast extract, as previously described (Silar 2020).All recovered isolates were heterokaryons with both *MAT1-1* and *MAT1-2* nuclei as all strains were self-fertile. Main characteristics of the strains are summarized in Table 1.

PSN309 F0 MAT1-1 and PSN309 F0 MAT1-2 were recovered by fragmenting a MAT*1-1*/MAT*1-2* heterokaryotic PSN309 F0 mycelium explant (Silar 2020). PSN1235_F1-sp1 and PSN1235_F1-sp3 were recovered by germinating small-sized ascospores expelled from perithecia obtained on PSN1235A cultures (Silar 2020). All strains are stored at -80°C in ESE and LIED labs and are available upon request.

#### DNA extraction and PCR amplification for ITS sequencing

For ITS sequencing, the genomic DNA was extracted using the rapid method previously described (Silar et al. 2025). The ITS locus was amplified using the ITS5 and ITS4 primers as recommended (White et al. 1990). The sequencing was performed by Genewiz (Azenta Life Science, Burlington, Massachusetts, U.S.A.), with primers ITS5 and ITS4. Final ITS sequences were manually assembled.

#### DNA extraction and short-read whole genome sequencing

The genome of the PSN309 F0 mycelium explant was previously sequenced by the JGI (Hensen et al. 2023). For Illumina genome sequencing of the other strains, the genomic DNA was extracted from fungal cultures grown on M2 Petri dishes using the NucleoSpin soil kit from Macherey-Nagel. The libraries used for sequencing and the Illumina sequencing were made by Novogene using a Illumina HiSeq2500 platform. The sequencing was performed using 2×150 bp Novaseq Illumina. The genomes were assembled using Unicycler version 0.5.1 with the default parameters (Wick et al. 2017) or Spades tool v3.11.1 (Bankevich et al. 2012). Assemblies for PSN824 and PSN1235_F1-sp1 were cleaned using the Foreign Contamination Screen (FCS-GX) tool from National Center for Biotechnology Information (NCBI)(Astashyn et al. 2024); all the other assemblies were exempt from foreign DNA contamination. The completeness of the genomes was evaluated by running the BUSCO program version 5.5.0 (Manni et al. 2021) on the Galaxy Europe server (https://usegalaxy.eu/ accessed on 26/03/2026) using the “Busco v5 lineage datasets (all+2024-03-21)”. The average nucleotide identities (ANI) were calculated using FungANI (Lalanne and Silar 2025) with the default parameters. To quantify the repeated sequences, a *de novo* identification of transposable elements was made with RepeatModeler v1.0.11 (Smit and Hubley 2008). RepeatMasker v4.0.9 (Smit et al. 2013) with the transposable elements detected by RepeatModeler as library source was subsequently used to quantify repeats. To predict the MAT idiomorph (MAT1-1 or MAT1-2), we performed tblastn analyses (Camacho et al. 2009) using the *P. anserina* MAT idiomorphs (*Pa_1_20590* for MAT1-2 and *Pa_1_20593, _20592, _20591* for MAT1_1). When one mating-type idiomorph was absent by blast in one genome, the genome was considered as homokaryotic for the following genomic analyses.

#### Phylogeny analyses

Sequences of the ITS + LSU sequences used to build the phylogenetic tree were extracted from GenBank (Table S2), the JGI website for *Cladorrhinum bulbillosum* DJ3 (https://mycocosm.jgi.doe.gov/Clabul1/Clabul1.home.html) and from genome sequences (Table S3). The concatenated sequences were aligned with MAFFT v7 (Katoh and Standley 2013) and the phylogenetic tree built using IQ-tree v2.4.0 (Minh et al. 2020). Statistical supports were calculated with 1000 ultrafast bootstrap replicates. A branch is considered well supported if its support is ≥0.95. For the phylogenomic-based tree, universal single-copy orthologs were identified using BUSCO v6 (Tegenfeldt et al. 2025) with the *sordariomycetes_odb12* lineage dataset. Only genes present as complete single-copy orthologs across all strains (see Table S3) were retained for downstream phylogenomic analyses, resulting in a dataset of 3,399 genes. For each BUSCO group, amino-acid sequences were aligned independently using MAFFT v7.526 (Katoh and Standley 2013) with the --auto option, allowing MAFFT to automatically select the most appropriate alignment strategy. The resulting alignments were trimmed using ClipKIT v2.12.0 (Steenwyk et al. 2020) with the kpic mode, retaining only parsimony-informative and constant sites. Filtered alignments were combined into a partitioned dataset (one partition per gene) and analyzed using IQ-TREE v3.1.2 (Wong et al. 2026). The best-fitting substitution model for each partition was selected using ModelFinder (option -m MFP), while accounting for heterogeneity among genes and sites using the --sampling GENESITE option. To reduce model overparameterization, similar partitions were merged using the relaxed clustering algorithm (option -rcluster 10). Phylogenetic inference was conducted using the maximum likelihood approach implemented in IQ-TREE. Branch support was assessed using 1,000 ultrafast bootstrap replicates.

### Long-read based genome sequencing, assembly and annotation

The homokaryotic *C. taenioides* PSN309 F0 MAT1-2 strain was additionally sequenced *in house* using the long-read sequencing Oxford Nanopore (MinION) technology as described in Vittorelli et al. 2023. Briefly, for DNA extraction, mycelium was cultured in liquid M2 medium during seven days and lyophilized. DNA was extracted with the NucleoBond High Molecular Weight DNA kit from Macherey-Nagel. The Nanopore library was prepared with the SQK-LSK109 ligation sequencing kit, and sequencing was performed with a R9.4.1 flow cell. The quality of Nanopore sequencing reads was assessed using PycoQC v2.5.2 (Leger and Leonardi 2019). MinION reads from the *C. taenioides* PSN309 F0 MAT1-2 strain were used to perform a *de novo* genome assembly using Flye v2.9.5-b1801 (Kolmogorov et al. 2019) with a genome size parameter set to 37 Mb (genomeSize=37m). This assembler performs read correction, filtering, and trimming of low-quality read ends. To correct assembly errors in the *C. taenioides* PSN309 F0 MAT1-2 genome, MinION reads were first mapped back to the assembly using Minimap2 v2.30 (Li 2018). The assembly was then polished through four iterative rounds of Racon v1.5.0 (Vaser et al. 2017) using parameters -m 8 -x -6 -g -8 -w 500, followed by consensus correction with Medaka v2.1.0 (Oxford Nanopore Technologies) according to the developer’s recommendations. For each polishing round, reads were realigned to the updated assembly using Minimap2. In a second step, Illumina short reads were used for additional polishing. Illumina reads were first aligned to the polished assembly using BWA v0.7.19 (Li 2013) and then, the alignment was used as input for Pilon v1.24 (Walker et al. 2014). Final assembly quality was assessed using QUAST v5.1 (Mikheenko et al. 2018) and BUSCO v6.0.0 (Tegenfeldt et al. 2025) with the *sordariomycetes_odb12* dataset in genome mode. BUSCO scores were used to estimate genome completeness. We refer to this assembly as the long-read-based *C. taenioides* PSN309 F0 MAT1-2 assembly. We checked that the mating-type chromosome assembly was identical when generated using an alternative assembly procedure described in Vittorelli et al. 2023.

Transposable elements were annotated using RepeatModeler v1.0.11 (Smit and Hubley 2008) to generate a *de novo* repeat library. RepeatMasker v4.0.9 (Smit et al. 2013) was used for annotation. Overlapping and fragmented annotations were merged using parseRM_merge_interrupted.pl (https://github.com/4ureliek/Parsing-RepeatMasker-Outputs; last accessed 31th May 2024). Elements shorter than 150 bp were removed to exclude low-complexity repeats. Gene prediction was then performed using the Helixer beta web server (Holst et al. 2023) with the “Fungi” model for both PSN309 F0 MAT1-1 and PSN309 F0 MAT1-2 genomes (last accessed 24th August 2026). Centromere positions were predicted using BLAST searches with centromere-flanking sequences of the mating-type chromosome from *P. anserina* (*Pa_1_12830*, *Pa_1_14290*, (Westerberg et al. 2025)). GC content was computed using the EMBOSS suite (Rice et al. 2000). The R package circlize was used to draw the circular plot in Figure S4C (Gu et al. 2014). Telomeric repeats were identified by screening contig ends for the canonical repeat motif TTAGGG and its reverse complement CCCTAA. Genome synteny was assessed using D-GENIES (Cabanettes and Klopp 2018), which relies on Minimap2 (Li 2018) alignments for visualization of genomic collinearity. To identify mitochondrial contigs, GC content variation across contigs was first examined using QUAST (Mikheenko et al. 2018) outputs, as mitochondrial genomes typically exhibit distinct GC content relative to nuclear DNA (Romiguier and Roux 2017). Putative mitochondrial contigs were further validated using BLAST v2.15.0 (Camacho et al. 2009) against the NCBI nucleotide database (nt).

### Mating-type locus annotation

To annotate MAT idiomorphs, translated BLAST searches (tBLASTn v2.15.0; (Camacho et al. 2009)) were performed using protein sequences of MAT-associated genes from *P. anserina*, called *FMR1* (=*mat1-1-1*), *SMR1* (=*mat1-1-2*), *SMR2* (=*mat1-1-3*) for the MAT1-1 idiomorph, and *FPR1* (=*mat1-2-1*) for the MAT1-2 idiomorph, as well as the flanking genes *APN2* and *SLA2*) (Debuchy and Coppin 1992) as queries in the long-read-based *C. taenioides* PSN309 F0 MAT1-2 assembly and the short-read-based *C. taenioides* PSN309 F0 MAT1-1 assembly. Gene boundaries were refined by manual inspection of start/stop codons and intron–exon junctions based on conserved splice-site motifs (Silar 2013) using the tool Artemis v18.1.0 (Carver et al. 2012). Final annotations are provided in Supplementary Fig 3.

### SNP calling

To assess the divergence between mating-type chromosomes, we performed SNP calling for five *Cladorrhinum* strains (PSN309, PSN1235, PSN672, PSN824 and PSN1024) against the long-read-based *C. taenioides* PSN309 F0 MAT1-2 assembly. We looked for SNP difference between the homokaryotic genomes PSN1235-sp1 and PSN1235-sp3,the homokaryotic genomes PSN1024C and PSN1024D for *C. neoaustrale* PSN1024, and the homokaryotic genomes PSN672C and PSN672D for *C. neoaustrale* PSN672. Missing data and heterozygote SNPs within each genomes were excluded. For *C. neoaustrale* PSN824, we looked for heterozygote SNPs within the heterokaryotic genome PSN824, excluding missing data. We first mapped Illumina reads using Bowtie2 v2.4.1 (Langmead et al. 2009), followed by duplicate removal using Picard MarkDuplicates v2.8.1 (Picard Toolkit, 2019). Variant calling was performed using GATK v4.1.2.0 (McKenna et al. 2010) with HaplotypeCaller. Variants were merged (CombineGVCFs), filtered (SelectVariants; SNP-only selection), and quality-filtered using VariantFiltration (QD < 20, FS > 60, MQ < 20, ReadPosRankSum ±2, MQRankSum ±2, SOR > 3, QUAL < 100). Filtered SNPs were retained using SelectVariants (--exclude-filtered). VCF files were converted to tabular format using VariantsToTable (-F CHROM -F POS -F TYPE -F REF -GF GT) and analysed in R v4.3.3 using RStudio v2023.12.1.402 and the tidyverse v2.0.0 suite. The heterozygous region surrounding the MAT locus was defined as the longest interval encompassing the MAT locus with its boundaries marked by the outermost SNPs such that no SNPs were more distant than 50 kb.

### Sexual crosses

All sexual crosses were performed on oatmeal agar (OA). MAT1-1 and MAT1-2 homokaryons, maintained on M2 medium, were subcultured onto OA Petri dishes. Each OA Petri dish received four plugs per strain (approximately 5 mm² each). The two parental strains were placed 1 cm apart on the same Petri dish. Plates were incubated at 27°C under continuous light. As the two mycelia grew and met, perithecia developed along the confrontation zone, producing ascospores after approximately 10–20 days. For heterokaryons, one plug was placed in the center of the Petri dish and perithecia developed on the growing mycelium. To collect ascospores from a cross, agar lids were placed above sporulating cultures to capture discharged ascospores. Individual spores were isolated under a stereomicroscope using sterile metal needles and plated on germination medium G. Spores were heat-shocked (60°C for 30 min followed by overnight incubation at 27°C or 36°C). The *C. taenioides* PSN1235 strain required a modified treatment consisting of incubation at 34°C overnight. Plates were then incubated at room temperature (∼20°C) during three to four days and subsequently stored at 4°C to halt growth.

To generate F1 spores of PSN309, two sexual crosses were performed between PSN309 F0 MAT1-2 and PSN309 F0 MAT1-1. Spores were arranged by tetrad (four ascospores per ascus) when possible. For the selfed progeny of PSN1235, crosses were performed between PSN1235-sp1 and PSN1235-sp3. For the selfed progeny of PSN1024, crosses were performed between PSN1024A and PSN1024C. For the selfed progeny of PSN672, sexual crosses were performed between PSN672C and PSN672D.

To generate F11 PSN672C progeny, ten generations of selfing were performed starting from the F1 PSN672C *C. neoaustrale* strain Each generation originated from a single heterokaryotic spore. For interspecies and intraspecies crosses, we used the parental strains described in Table S4.

### Genotyping of the F11 PSN672C progeny

We sequenced genomes of the mycelium growing from 15 F11 PSN672C heterokaryotic spores as described above in “DNA extraction and short-read whole genome sequencing”. To study heterozygosity around the MAT locus, we called SNPs as described above in “SNP calling”.

### Genotyping of the F1 PSN309 progeny

Genomic DNA was extracted from germinated spores using a rapid Chelex-based protocol (Vittorelli et al. 2023). PCR amplification of mating-type idiomorphs was performed using primers designed with Primer3 (Untergasser et al. 2012) and validated by BLAST (Camacho et al. 2009)(Table S8). One spore per ascus was initially genotyped, and a subset of asci was fully dissected for detailed segregation analysis. Spores and full asci were isolated after projection on agar medium. PCR was performed following conditions described in Table S9. PCR products were visualized on 1.5% agarose gels containing GelRed in 0.5× TAE buffer, run at 135 V for 30 minutes.

### Self-fertility assays of single-spore-derived mycelia

For self-fertility assessment, all germinated spores were transferred onto OA medium five days after isolation. For the *C. australis* PSN672C strain, the assay was first performed both on 101 F11 PSN672C spores and 25 F1 PSN672C spores. For all other strains, analyses were conducted on F1 spores, corresponding to the products of a single sexual generation of selfing from the isolated strains from nature, with sample sizes ranging from 18 to 30 spores depending on the strain.

To estimate the proportion of heterokaryotic spores, germinated spores were transferred individually onto OA medium for *Cladorrhinum* strains. Plates were incubated at 22°C under continuous light. Mycelial growth was monitored for several weeks, and the appearance of perithecia and discharged ascospores was recorded. Single-spore-derived mycelia capable of producing perithecia and ascospores were scored as self-fertile. Because self-fertility requires the coexistence of nuclei of opposite mating types, the corresponding spores were inferred to be heterokaryotic.

Statistics tests were performed with the software JUMP SAS Institute (Version 3.1.5, JMP Statistical Discovery LLC, Cary, NC). Plots were performed with R ggplot2 software (Wickham 2009).

## Data Availability Statement

All newly generated genome data were deposited on the NCBI Genbank platform under the BioProject ID number under request (see details in Table S1).

## Author contribution

T.G., F.E.H., and P.S. conceptualized and supervised the study. T.G., F.E.H., and P.S. acquired funding. T.G. and P.S. collected the samples. V.G. and P.S. isolated the strains. E.C., V.G and J.L.-D. performed experiments of culture in vitro and molecular biology. F.E.H, C.L., J.L.-D., and P.S. analyzed the genomes. E.C., F.E.H., J.L.-D, and P.S. produced the figures. T.G., F.E.H., J.L.-D, and P.S. wrote the original draft. All authors edited the manuscript.

## Acknowledgments

We thank Gabriela Rezende, Elizabeth Chahine and Amandine Labat for help in DNA extraction, sequencing and mycelium culturing experiments, Alodie Snirc for help in the laboratory and Ricardo Rodriguez de la Vega for help with bioinformatic analyses. We thank Jean-Philippe Vernadet for help in formatting and publishing genomic data. We thank Lou Guyot for discussing ideas and for comments on previous versions of the manuscript. We thank all sample collectors: Roselyne Ferrari, Pierre Defos du Rau, Yves Hurand, and Lucas Bonometti. This work was supported by the Louis D. Foundation award and ERC advanced grants EvolSexChrom #832352 and DEGENSEX #101262718 to TG.

