## Supplementary material for "Recombination persists near the mating-type locus in pseudo-homothallic *Cladorrhinum* fungi (Sordariales)"

**Supplementary Tables:**

**-Table S1: Main features of the Illumina-based genome assemblies: strain ID, culture type, sequencing technology, assembly method and size, contig number, contig N50 (sequence length such that 50% of the total assembly length is contained in contigs of that length or longer, in kilobases kb), size of the largest contig, of the unique regions and of the repeated regions (in Megabases Mb), BUSCO score (Benchmarking Universal Single-Copy Orthologs), mating type found in the assembly, and reference of the genome assembly.** Assemblies used for average nucleotide identity (ANI) calculations are indicated by an an asterisk *.

**-Table S2: GenBank accession numbers of the sequences used in the ITS + LSU tree.**

**-Table S3: Strain ID and taxonomy, and genome accession numbers of the assemblies used for phylogenomic analysis.**

**-Table S4: Results of inter- and intra-specific crosses of *Cladorrhinum taenioides* and *C. neoaustrale*.**

**-Table S5: List of predicted protein functions for 19 genes annotated in the region of heterozygosity around the mating-type locus in the *Cladorrhinum taenioides* PSN309 F0 MAT-1-2 strain.** Gene ontology terms predicted for proteins of the PSN309 F0 genome (Hensen et al. 2023) are presented. Correspondence of the gene id in the long read based assembly of PSN309 F0 MAT-1-2 strain and the PSN309 F0 genome assembly (Hensen et al. 2023) are indicated.

**-Table S6: Statistics of the heterozygous region around the mating-type locus in the studied *Cladorrhinum taenioides* and *C. neoaustrale* strains.** The start and end positions of the region on contig C_1 (bp), the size of the region (in bp) and the number of heterozygous SNPs are presented. In addition, for the F11 progeny of *Cladorrhinum neoaustrale* PSN672, accession ID and BioProject ID of each genome are presented.

**-Table S7: Percentages of ascospore-derived mycelia that were found heterokaryotic based on tests of mycelium self-fertility in selfed progeny of *Cladorrhinum taenioides* and *C. neoaustrale* strains.**

**-Table S8: Primer sequence used to genotype the MAT1_1 and MAT1_2 mating-type idiomorphs.**

**-Table S9: PCR conditions used to genotype the mating-type locus.**

**Figure S1. Segregation of the mating-type locus during meiosis in the closely related model species *Podospora anserina* and hypotheses for the loss of one mating type in mycelia.**
**A.** Second meiotic division (SDS) at the mating-type locus leading to the formation of four heterokaryotic/dikaryotic spores carrying both mating types. Only the chromosomes carrying the mating-type locus are represented, with their centromeres at the diploid and meiosis I stages (adapted from (Vittorelli et al. 2023), schema using biorender).
**B.** Hypotheses for the detection of a single mating type during mycelium genotyping despite SDS at the mating-type locus. (1) Loss of nuclei during spore development leading to a multinucleate homokaryotic spore and homokaryotic mycelium; (2) Formation of heterokaryotic spores with equilibria between nuclei of opposite mating types, followed by the loss of nuclei during mycelium growth, sometimes only in specific sectors; (3) Disequilibria between nuclei of opposite mating types in heterokaryotic spores leading to a chimeric mycelium with homokaryotic and heterokaryotic sectors.

**Figure S2. Example of results for one interspecies cross of *Cladorrhinum taenioides* and *C. neoaustrale* and one intraspecific cross of *C. neoaustrale*. A.** Example of an interspecific cross: *C. taenioides* PSN309 F0 MAT1-2 x *C. neoaustrale* PSN672D MAT1-1 ; **B.** Example of an intraspecific cross : *C. neoaustrale* PSN1024D MAT1-2 x *C. neoaustrale* PSN672D MAT1-1. (1) Overview of the Petri dish. (2) A zoom was performed using binocular loop (x0.8) to show perithecia at the frontier of parental thalli. (3) Squashed perithecia and ascospores content as observed using binocular loop (x 5.6).

**Figure S3. Percentage of genome base pairs covered by transposable element families in the *Cladorrhinum taenioides* PSN309 F0 MAT1-2 long-read-based genome assembly.** LTR: Long Terminal Repeat retrotransposons; LINE : Long Interspersed Nuclear Elements retrotransposons. DNA/TcMar : DNA transposon Tc1/mariner**.**

**Figure S4. Genomic characteristics of the putative centromere region of the mating-type chromosome (contig C_1) in the *Cladorrhinum taenioides* PSN309 F0 MAT1-2 long-read-based genome assembly.** A-B. GC content (panel A) and transposable element density (panel B) in percentages per 1 kb non-overlapping genomic region. The position of the putative centromere region (4,233,487-4,242,272) is represented by a vertical dash green line. C. Synteny between the mating-type chromosomes of *C. taenioides* PSN309 F0 MAT1-2 (purple, bottom) and *Podospora anserina* S strain (blue, top), that is closely related. The non recombining region is indicated in red (corresponding to the genomic region that is heterozygous between mating types) and the putative centromere region in yellow. Orthologous genes in single copy are indicated by the links in color.

**Figure S5. Mating-type locus annotation and synteny in the MAT1-1 and MAT1-2 genomes of *Cladorrhinum taenioides* PSN309. A-B.** The mating-type locus in the Illumina-based assembly of *C. taenioides* PSN309 F0 MAT1-1 (A) and the long-read based assembly of *C. taenioides* PSN309 F0 MAT1-2 (B) assemblies viewed in Artemis v18.0.2 (Carver et al. 2012). The *APN2* and *SLA2* genes flanking the mating-type locus are indicated. The green boxes represent the genomic regions differing between MAT1-1 and MAT1-2 genomes. The three forward and three reverse frames of the DNA sequence are presented; vertical bars indicate putative STOP codons. C. Synteny in the mating-type locus region as viewed in D-GENIES (Cabanettes and Klopp 2018) between the Illumina-based assembly of *C. taenioides* PSN309 F0 MAT1-1 (contigs > 27 kb are shown) and the long-read based assembly of *C. taenioides* PSN309 F0 MAT1-2 The region of 5.7, 6.4 Mb of the C_1 contig is shown.

**Figure S6. Genome-wide heterozygosity in several *Cladorrhinum* strains.** A. Heterozygous SNPs per 50-kb windows in *C. taenioides* PSN1235 strain, as assessed by SNPs differences between the homokaryotic genomes PSN1235-sp1 and PSN1235-sp3. B. Heterozygous SNPs per 50-kb windows in the heterokaryotic genome of *C. neoaustrale* PSN824; C. Heterozygous SNPs per 50-kb windows in *C. neoaustrale* PSN1024 strain, as assessed by SNPs differences between the homokaryotic genomes PSN1024C and PSN1024D. D. Heterozygous SNPs per 50-kb windows in *C. neoaustrale* PSN672 strain, as assessed by SNPs differences between the homokaryotic genomes PSN672C and PSN672D.

**Figure S7. Reduction in heterozygosity around the mating-type locus in the F11 progeny of PSN672C *Cladorrhinum neoaustrale* strain.** Genome-wide distribution of heterozygous SNPs per 50 kb windows in the heterokaryotic genome of the F11 progeny PSN672 sp116.

**Figure S8. Intra-strain crosses for *Cladorrhinum taenioides* PSN309 and PSN1235 (A) and *C. neoaustrale* PSN672 and PSN1024 (B).** Overview of the Petri dish. **A.** (1) *C. taenioides* PSN309 F0 MAT1-2 x PSN309 F0 MAT1-1; (2) *C. taenioides* PSN1235_F1-sp1 MAT1-2 x PSN1235_F1-sp3 MAT1-1; **B.** (1) *C. neoaustrale* PSN672C MAT1-2-x PSN672D MAT1-1; (2) *C. neoaustrale* PSN1024D MAT1-2 x PSN1024A MAT1-1.

Figure S1

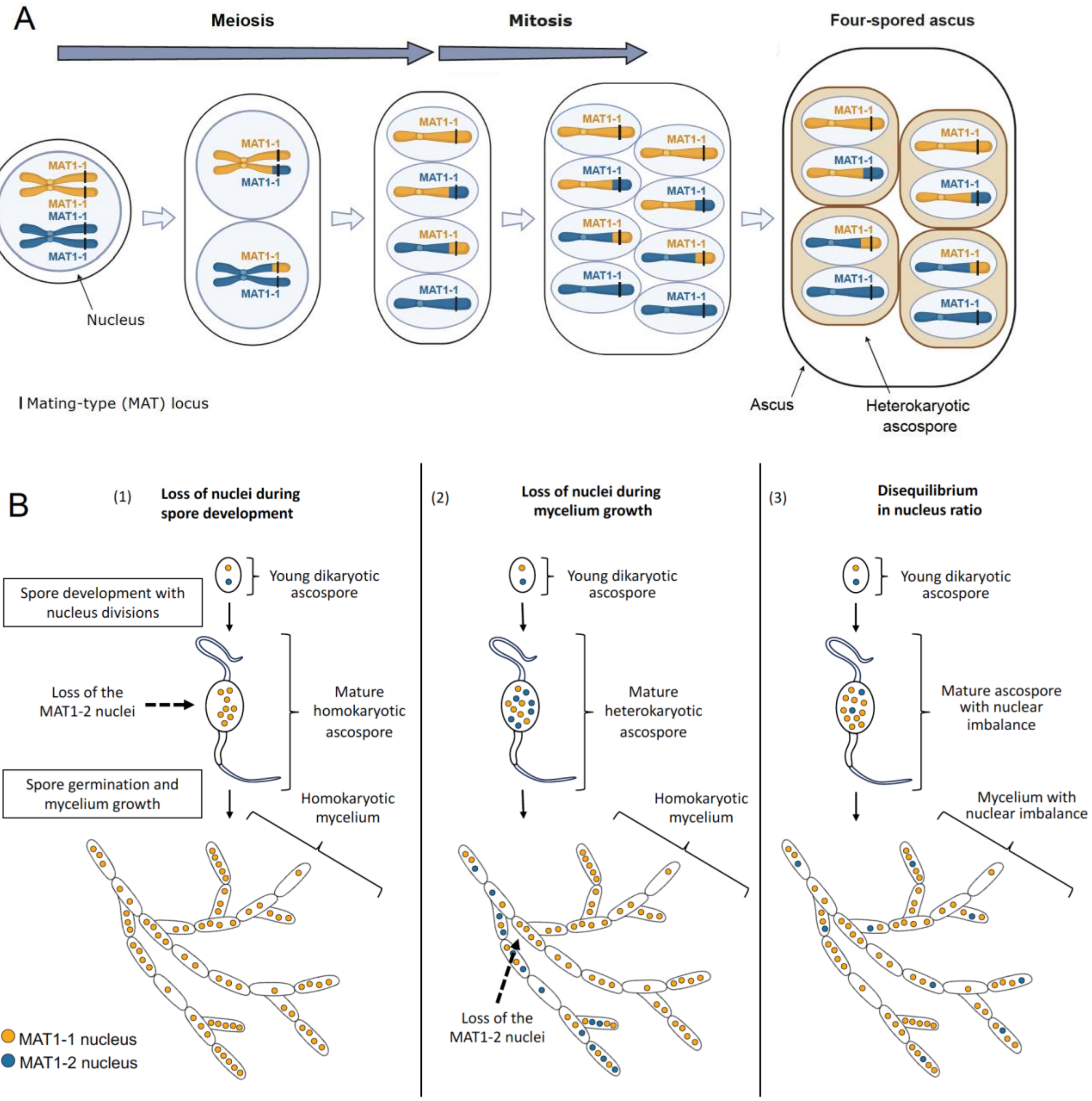

A
Meiosis
Mitosis
Four-spored ascus
MAT1-1
Nucleus
Mating-type (MAT) locus
Ascus
Heterokaryotic ascospore
B
(1) Loss of nuclei during spore development
(2) Loss of nuclei during mycelium growth
(3) Disequilibrium in nucleus ratio
Young dikaryotic ascospore
Spore development with nucleus divisions
Loss of the MAT1-2 nuclei
Mature homokaryotic ascospore
Spore germination and mycelium growth
Homokaryotic mycelium
Mature heterokaryotic ascospore
Mature ascospore with nuclear imbalance
Mycelium with nuclear imbalance
MAT1-1 nucleus
MAT1-2 nucleus

Figure S2

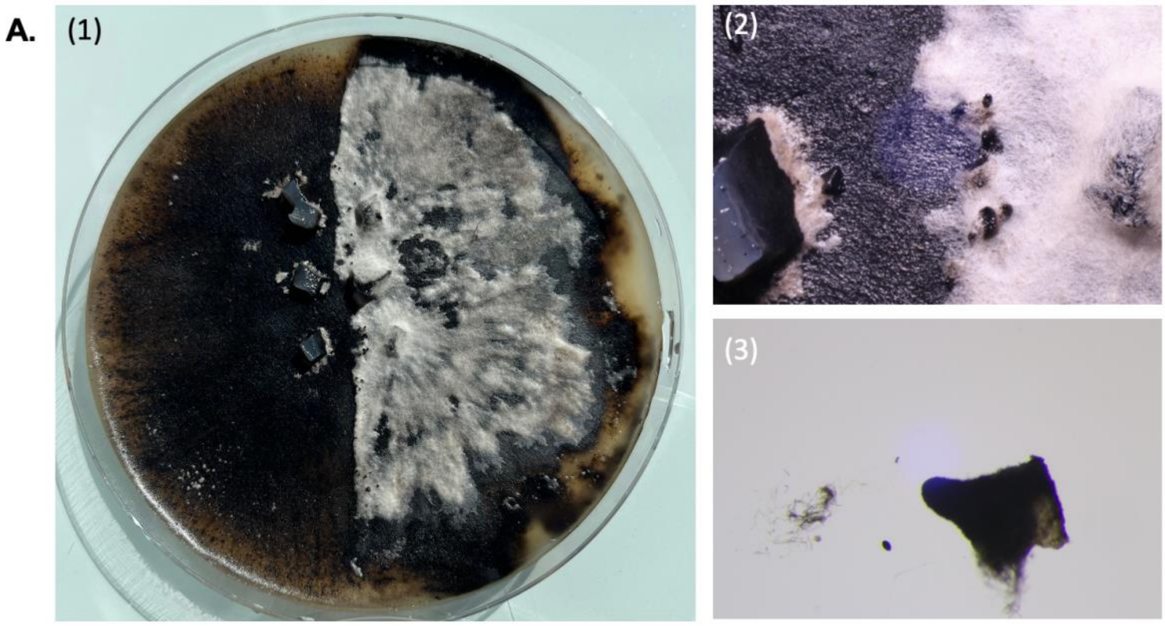


PSN309_F0-mat1-2 x PSN672D mat1-1

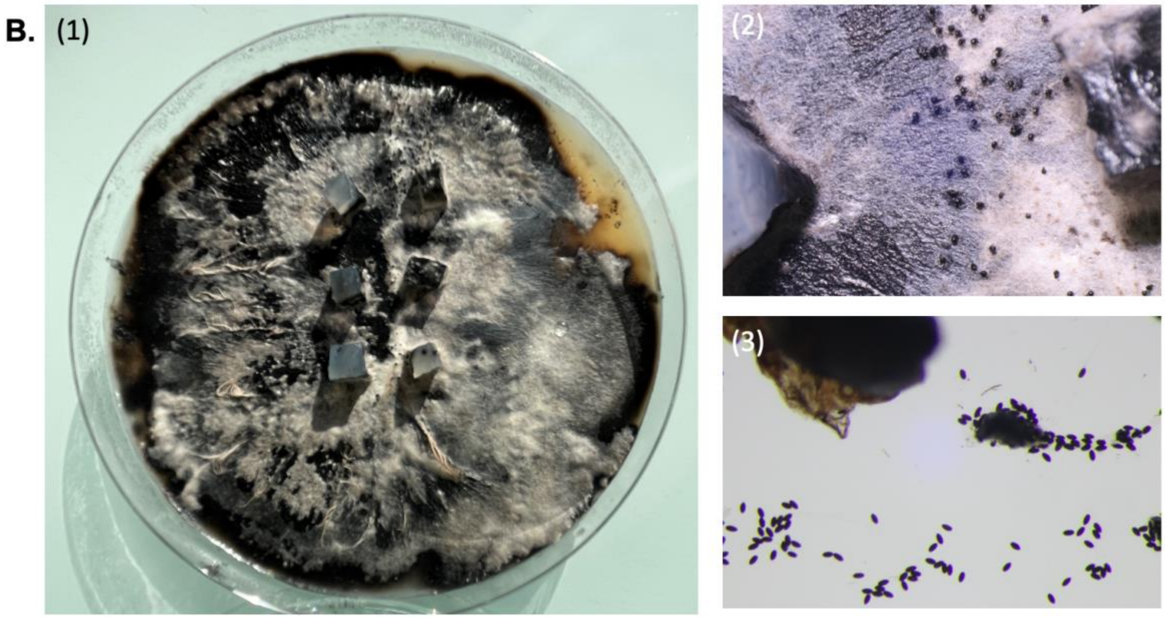


PSN1024D mat1-2 x PSN672D mat1-1

Figure S3

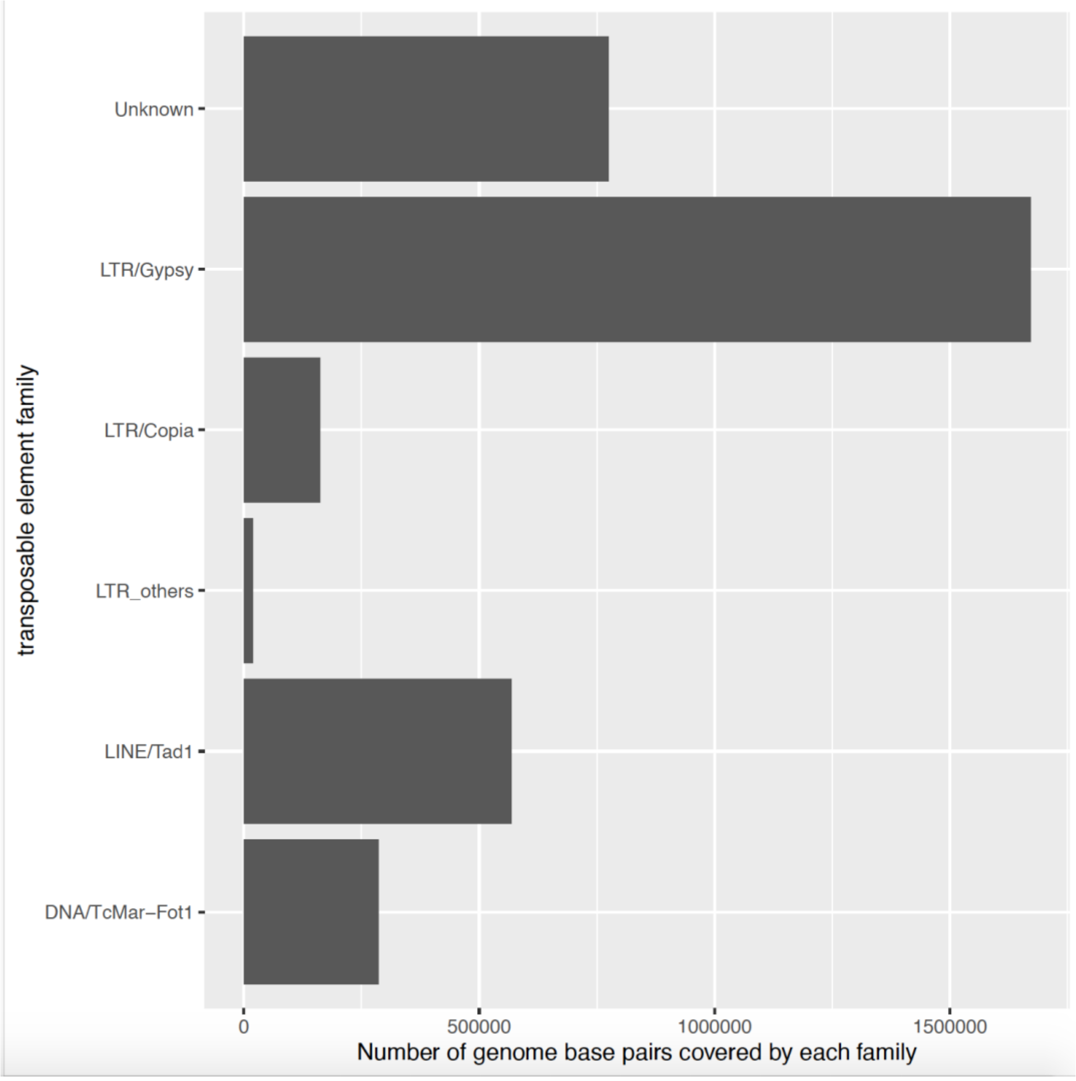

transposable element family
Unknown
LTR/Gypsy
LTR/Copia
LTR_others
LINE/Tad1
DNA/TcMar–Fot1
0
500000
1000000
1500000
Number of genome base pairs covered by each family

Figure S4

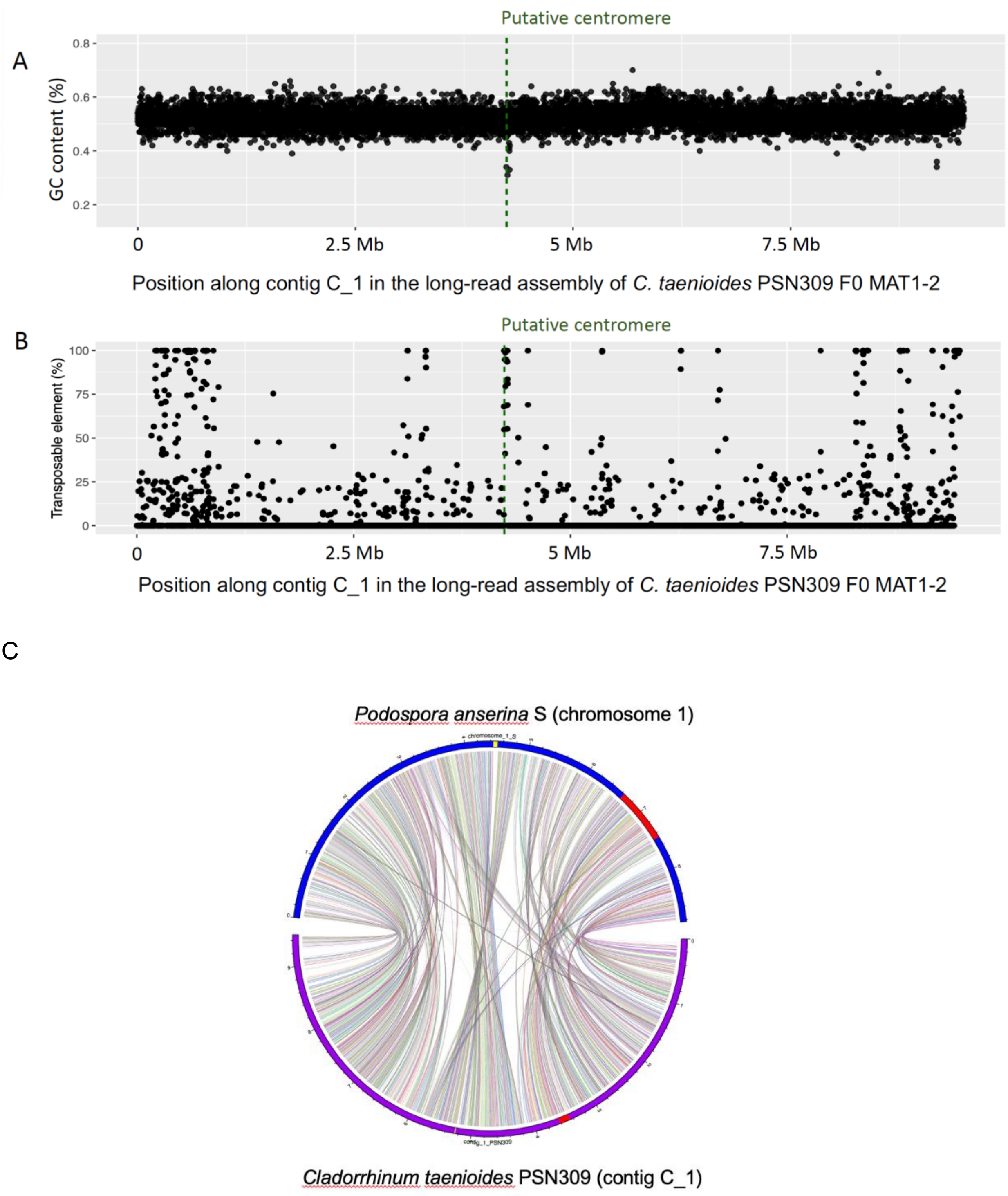
A
Putative centromere
GC content (%)
0.8
0.6
0.4
0.2
0
2.5 Mb
5 Mb
7.5 Mb
Position along contig C_1 in the long-read assembly of *C. taenioides* PSN309 F0 MAT1-2
B
Putative centromere
Transposable element (%)
100
75
50
25
0
0
2.5 Mb
5 Mb
7.5 Mb
Position along contig C_1 in the long-read assembly of *C. taenioides* PSN309 F0 MAT1-2
C
*Podospora anserina* S (chromosome 1)
*Cladorrhinum taenioides* PSN309 (contig C_1)

Figure S5 :

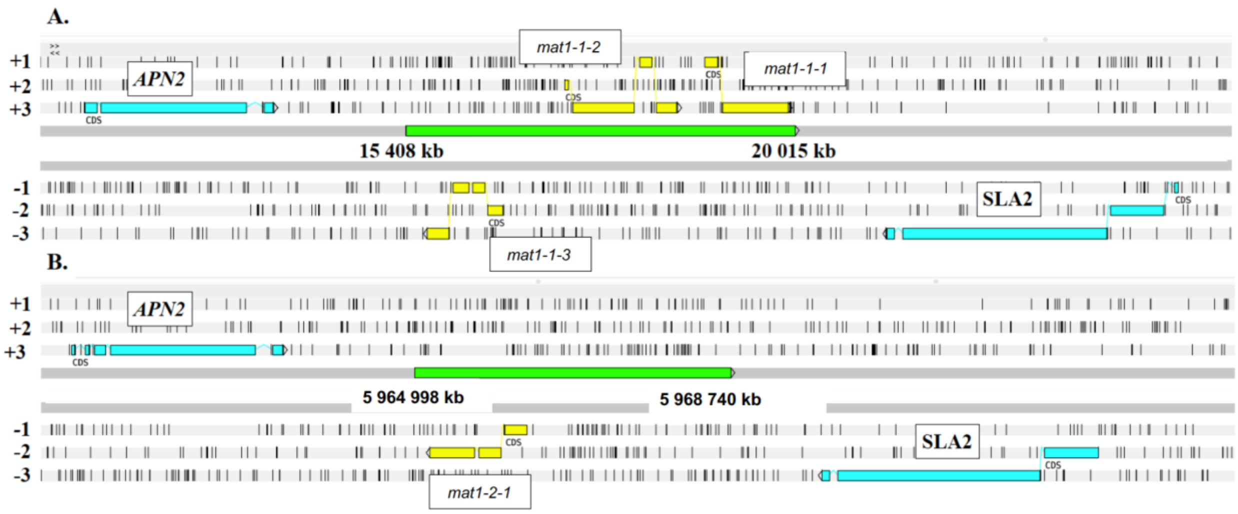
A.
+1
+2
+3
APN2
mat1-1-2
mat1-1-1
15 408 kb
20 015 kb
-1
-2
-3
SLA2
mat1-1-3
B.
+1
+2
+3
APN2
5 964 998 kb
5 968 740 kb
-1
-2
-3
SLA2
mat1-2-1

C

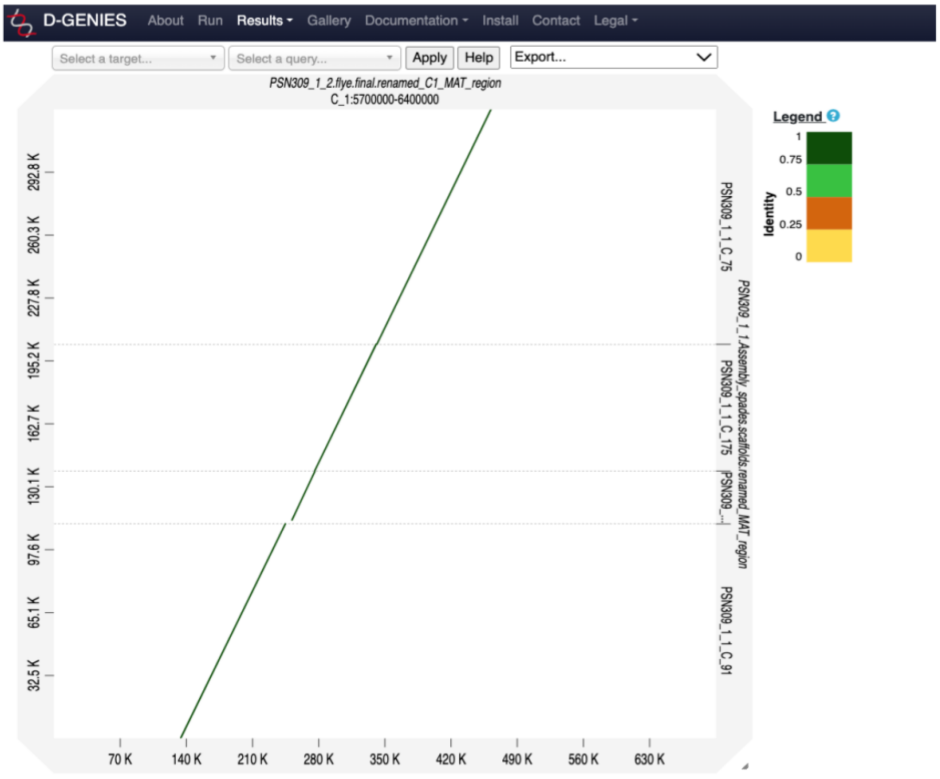
D-GENIES
About
Run
Results
Gallery
Documentation
Install
Contact
Legal
Select a target...
Select a query...
Apply
Help
Export...
PSN309_1_2.flye.final.renamed_C1_MAT_region
C_1:5700000-6400000
Legend
Identity
1
0.75
0.5
0.25
0
292.8 K
260.3 K
227.8 K
195.2 K
162.7 K
130.1 K
97.6 K
65.1 K
32.5 K
70 K
140 K
210 K
280 K
350 K
420 K
490 K
560 K
630 K
PSN309_1_1_C_75
PSN309_1_1_C_175
PSN309_1_1_C_91
PSN309_1_1.Assembly_spades.scaffolds.renamed_MAT_region

Figure S6:

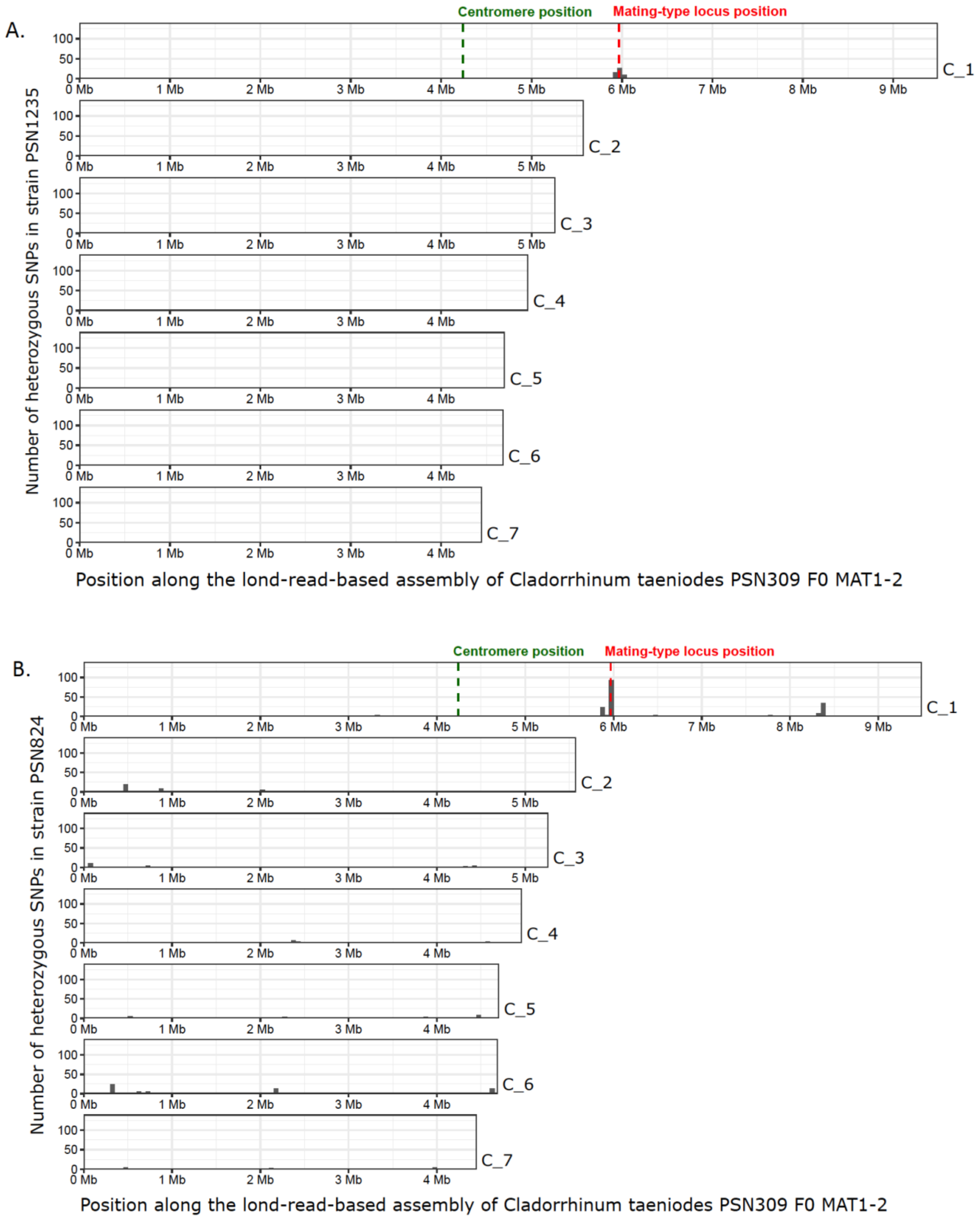

A.
Centromere position
Mating-type locus position
Number of heterozygous SNPs in strain PSN1235
C_1
C_2
C_3
C_4
C_5
C_6
C_7
Position along the lond-read-based assembly of Cladorrhinum taeniodes PSN309 F0 MAT1-2
B.
Centromere position
Mating-type locus position
Number of heterozygous SNPs in strain PSN824
C_1
C_2
C_3
C_4
C_5
C_6
C_7
Position along the lond-read-based assembly of Cladorrhinum taeniodes PSN309 F0 MAT1-2

C.

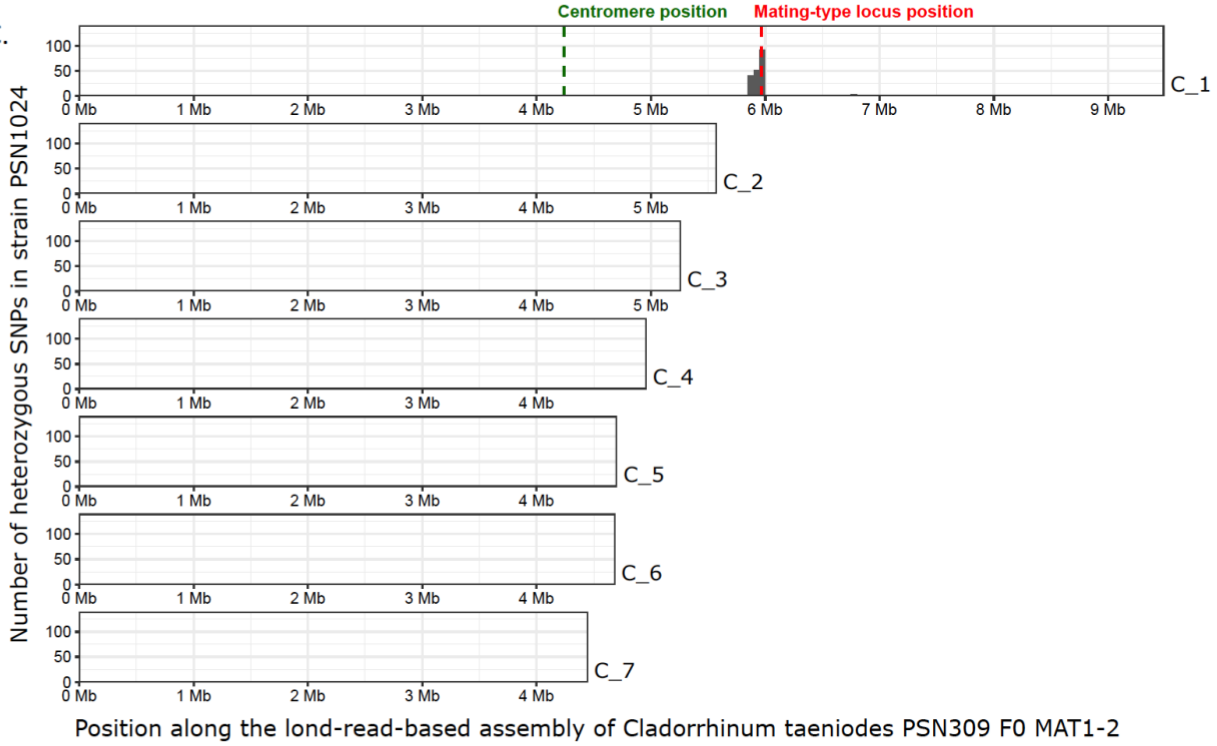
Centromere position
Mating-type locus position
Number of heterozygous SNPs in strain PSN1024
100
50
0
0 Mb
1 Mb
2 Mb
3 Mb
4 Mb
5 Mb
6 Mb
7 Mb
8 Mb
9 Mb
C_1
C_2
C_3
C_4
C_5
C_6
C_7
Position along the lond-read-based assembly of Cladorrhinum taeniodes PSN309 F0 MAT1-2

D.

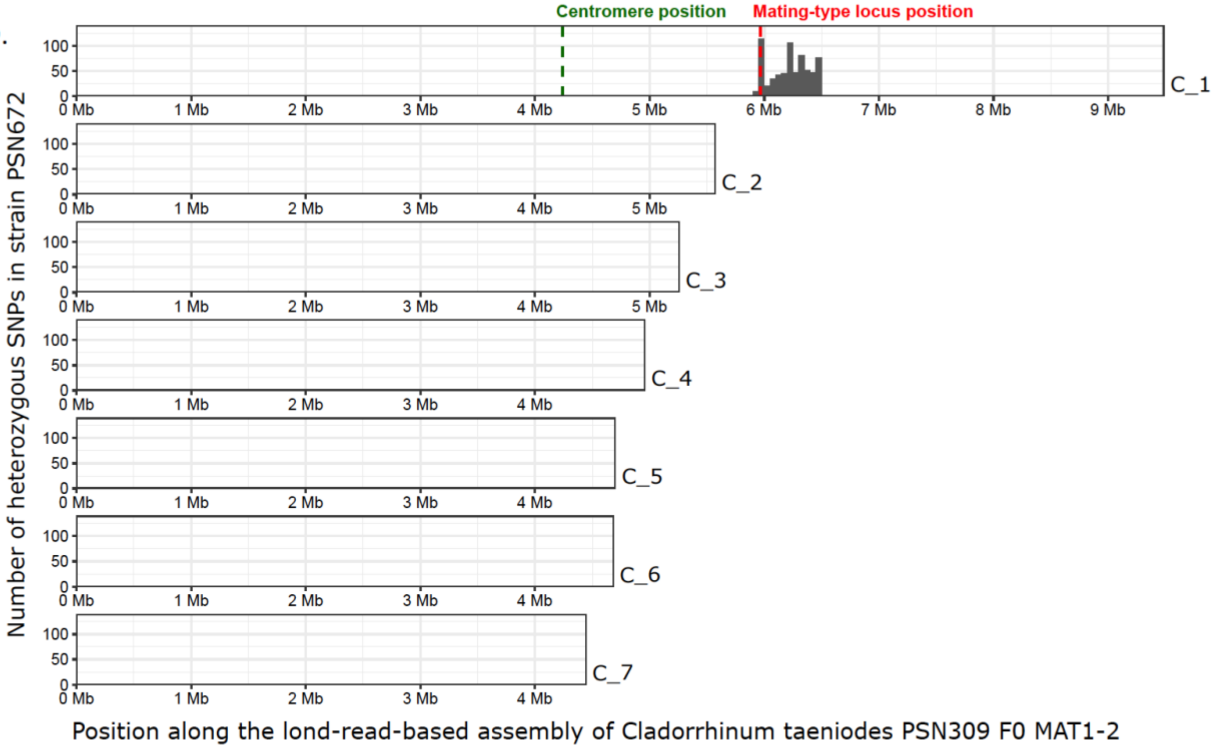
Centromere position
Mating-type locus position
Number of heterozygous SNPs in strain PSN672
100
50
0
0 Mb
1 Mb
2 Mb
3 Mb
4 Mb
5 Mb
6 Mb
7 Mb
8 Mb
9 Mb
C_1
C_2
C_3
C_4
C_5
C_6
C_7
Position along the lond-read-based assembly of Cladorrhinum taeniodes PSN309 F0 MAT1-2

Figure S7

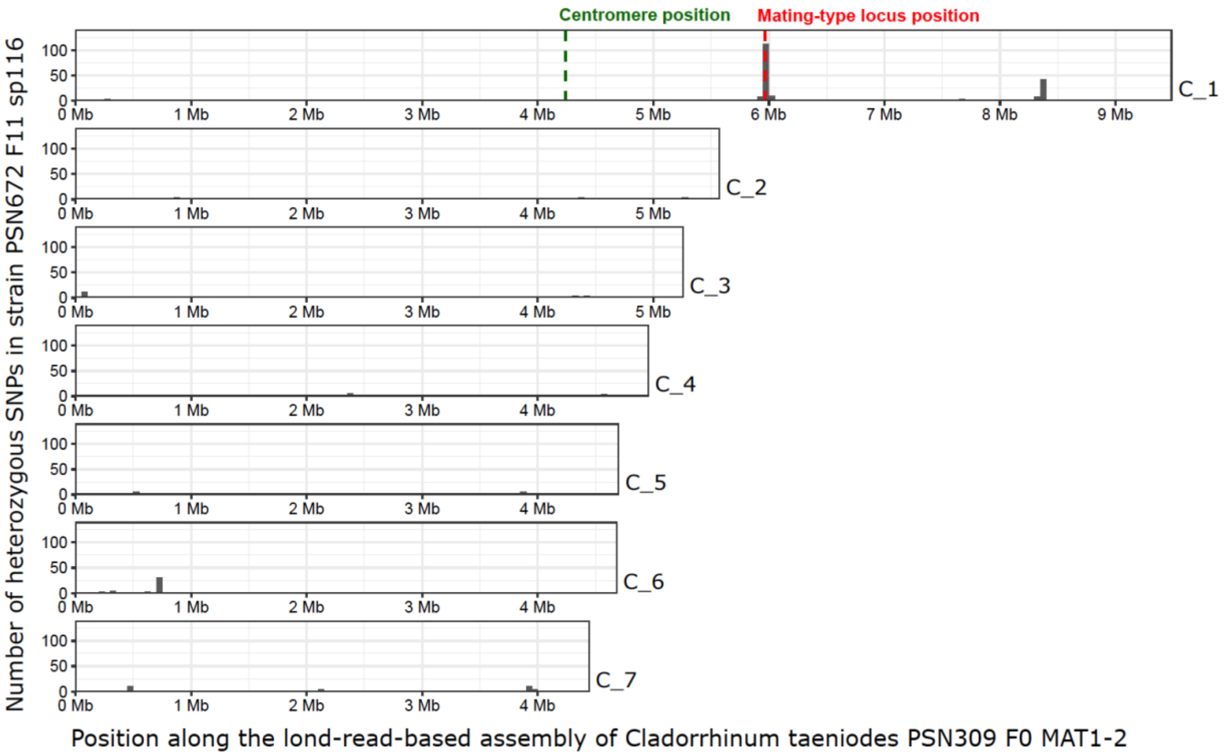

Centromere position
Mating-type locus position
Number of heterozygous SNPs in strain PSN672 F11 sp116
100
50
0
0 Mb
1 Mb
2 Mb
3 Mb
4 Mb
5 Mb
6 Mb
7 Mb
8 Mb
9 Mb
C_1
C_2
C_3
C_4
C_5
C_6
C_7
Position along the lond-read-based assembly of Cladorrhinum taeniodes PSN309 F0 MAT1-2

Figure S8

**A.** (1)

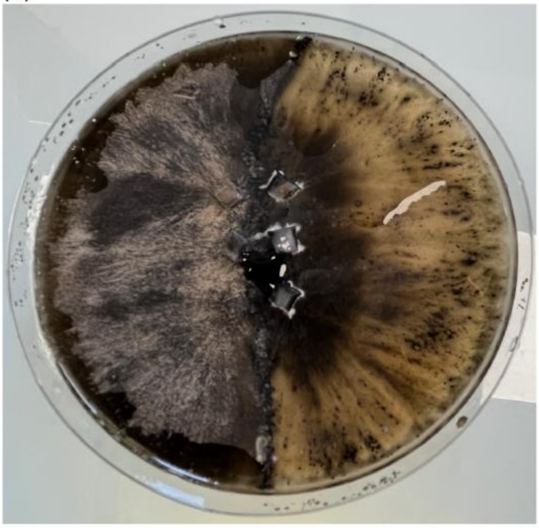

PSN309_F0-mat1-2 x PSN309_F0-mat1-1

(2)

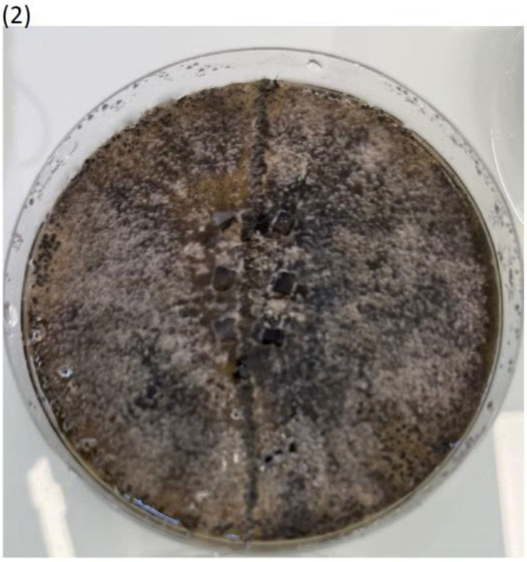

PSN1235_F1-sp1-mat1-2--x-PSN1235_F1-sp3-mat1-1

**B.** (1)

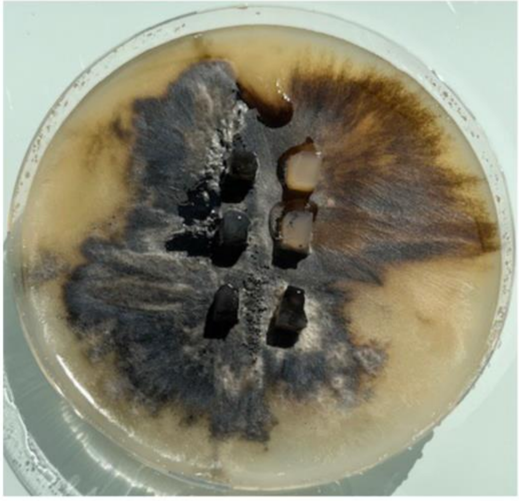

PSN672C-mat1-2-x-PSN672D-mat1-1

(2)

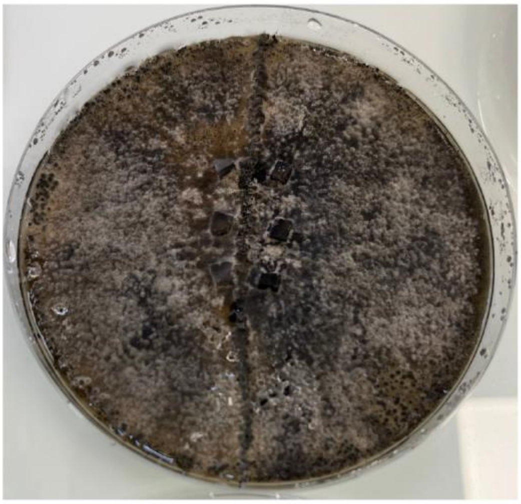

PSN1024D-mat1-2-x-PSN1024A-mat1-1
